# Macrofaunal Diversity and Community Structure of the DeSoto Canyon and Adjacent Slope

**DOI:** 10.1101/2020.01.15.908194

**Authors:** Arvind K. Shantharam, Chih-Lin Wei, Mauricio Silva, Amy R. Baco

**Affiliations:** Department of Earth, Ocean, and Atmospheric Sciences, Florida State University, 1011 Academic Way, Tallahassee, FL 32306; Institute of Oceanography, National Taiwan University, Taipei 106, Taiwan

## Abstract

Macrofauna within the DeSoto Canyon, northern Gulf of Mexico (GOM), along the canyon wall and axis, and on the adjacent slope, were sampled along with sediment, terrain, and water mass parameters. Within the canyon, abundance and species richness decreased with depth, while evenness increased. Cluster analysis identified three depth-related groups within the canyon that conformed to previously established bathymetric boundaries: stations at 464 – 485 m, 669 – 1834 m, and > 2000 m. Abundance differed between depth groups. Species richness was lowest for the deepest group and evenness was lowest for the shallowest. Community structure within the canyon most related to fluorometry and oxygen saturation, combined with any of salinity, particulate organic carbon, sediment organic carbon, or slope.

Canyon wall abundances were higher than the canyon axis or adjacent slope. Community structure differed between all three habitat types. Ordination of community structure suggests a longitudinal pattern that potentially tracks with increasing sea-surface chlorophyll that occurs in the eastward direction across the northern GOM. Canyon and slope differences may result from seasonal water masses entrained by canyon topography characterized by high salinity, oxygen saturation, fluorometry, and turbidity. Higher fluorescence and turbidity in the canyon did not translate into higher sediment organic matter. Flushing along canyon wall channels and the canyon axis may explain the low organic matter. Differences in abundance and structure between the canyon wall and axis may result from microhabitat heterogeneity due to potential hydrocarbon seepage, organically enriched sediment deposits along channels, or remnant influence from the Deepwater Horizon blowout.

## 1. Introduction

Submarine canyons are one of the most common large-scale bathymetric features in oceanic basins around the world (Harris & Whiteway 2011). Over 9540 have been detected along continental margins (Harris et al. 2014). They are known as hotspots of benthic biodiversity and biomass, receiving increasing attention from deep-sea researchers (Rowe et al. 1982, Houston & Haedrich 1984, Gerino et al. 1995, Maurer et al. 1995, Vetter & Dayton 1998, Sorbe 1999, Curdia et al. 2004, Tyler et al. 2009, De Leo et al. 2010, McClain & Barry 2010, Cunha et al. 2011a, Paterson et al. 2011, Hunter et al. 2013, Gunton et al. 2015, Harriague et al. 2019). Through an interplay of local hydrography and canyon topography, canyons may channel currents and format upwelling (Klinck 1996, Hickey 1997, Canals et al. 2006), entrain particulate organic matter (Vetter 1994, Vetter & Dayton 1998, Harrold et al. 2003, Company et al. 2008, Rowe et al. 2008, De Leo et al. 2010, De Leo et al. 2012, Hunter et al. 2013), and transport shelf sediments to slopes in episodic turbidity currents or mass-wasting events (de Stigter et al. 2007, Oliveira et al. 2007, Arzola et al. 2008). This, in turn, concentrates diel vertical migrators (Greene et al. 1988, Lavoie et al. 2000, Genin 2004), and provides enhanced seafloor habitat heterogeneity (Yoklavich et al. 2000, Brodeur 2001, Uiblein et al. 2003, Vetter et al. 2010, De Leo et al. 2012).

High seafloor habitat heterogeneity in turn enhances canyon benthic biodiversity. It can create a patchwork availability of resources that result in gradients in density and faunal turnover on 1 m – 1 km spatial scales (McClain & Barry 2010, De Leo et al. 2014, Campanyà-Llovet et al. 2018). Entrained particulate organic matter accumulates and shifts in distribution within a canyon to structure faunal density, biodiversity, biomass, and community structure (Vetter & Dayton 1999, Curdia et al. 2004, Escobar-Briones et al. 2008). Topographically-induced hydrographic and biochemical regimes are important sources of continual disturbance and have been noted to elevate abundance, biomass, and species richness compared to adjacent non-canyon regions (Duineveld et al. 2001, De Leo et al. 2010, De Leo et al. 2014, Harriague et al. 2019). All of this contributes to canyon denizens constituting a large proportion of marine metazoan benthic biodiversity and production (Gage 1996, Snelgrove 1999, Ebbe et al. 2010).

Submarine canyons located in the Gulf of Mexico (GOM) have received minimal attention in terms of the environmental and habitat heterogeneity and the effect these have on resident benthic macrofauna. What has been done has shown that major GOM depressions and canyons have high abundance and biomass, which has primarily been linked to large-scale processes such as the Mississippi River outflow, particulate organic matter flux, or grain (Baguley et al. 2006a, Baguley et al. 2006b, Escobar-Briones et al. 2008, Wei et al. 2012, Wei & Rowe 2019). This, however, does not account for more local scale processes and microhabitats that could strongly influence ecological processes affecting the macrobenthos.

The DeSoto Canyon, in the northeastern GOM, has been noted to contain high benthic decapod diversity (Wicksten & Packard 2005) and high abundances of infaunal organisms including both meiofauna (Baguley et al. 2006a) and macrofauna (Wei et al. 2010) compared to adjacent GOM sites. Macrofaunal biomass is also significantly higher in the canyon (Wei et al. 2012) and has been attributed to the high amount of particulate organic carbon (POC) entrained there (Morse & Beazley 2008, Wei & Rowe 2019) and perhaps from the large amount of continental shelf export it receives (Hamilton et al. 2015). Highly productive habitats such as hydrocarbon seeps also occur in the canyon (Washburn et al. 2018). While the question of how benthic species richness differs in the canyon compared to the slope has been addressed (Wei et al. 2019), a comparison of community structure largely has not.

The only previous study considering macrofaunal community structure of the DeSoto canyon was part of larger comprehensive investigations of the northern Gulf of Mexico (NGOM) macrobenthos, with a only a few samples collected in the canyon (Wei et al. 2010). Differences in canyon and slope community structure was not explicitly tested but comparisons can be inferred from the available data. The shallower of the canyon sites in that study, labeled S35, and a deeper canyon site, S36 clustered with non-canyon sites to the east and west, and were grouped into an “eastern mid-slope” zone. Similarly the deepest site in the canyon also clustered with non-canyon sites in its depth range (Wei et al 2010). These results suggest no difference within the canyon across a depth range of 1,721 m, nor between communities within the canyon compared to non-canyons sites. However, marine canyons are known to exhibit high amounts of beta diversity over small and large spatial scales for organisms in macrofaunal and megafaunal size classes (Schlacher et al. 2007, McClain & Barry 2010, Campanyà-Llovet et al. 2018). For example, Schlacher et al. (2007) reported highly restricted megafaunal sponge distributions in southeastern Australian canyons with 76% of species occupying a single site and 79% inhabiting single canyons. McClain & Barry (2010) observed high macrobenthic turnover (∼40%) between open canyon sites and sites closer (< 100 m) to the cliff faces of Monterey Canyon. On even smaller spatial scales, 10s of m apart, Campanyà-Llovet et al. (2018) found distinct macrofauna communities in the Barkley Canyon.

These studies suggest a potential for significant spatial variability in macrofaunal communities within the DeSoto Canyon that may have been missed at the coarse sampling scales previously undertaken. Therefore, the goal of this study was to characterize finer-scale spatial variability in the DeSoto Canyon macrofauna, particularly among the canyon wall, axis, and adjacent slope, across a range of depths, by testing for differences between macrofaunal abundance, diversity, and community structure (1) within the canyon, (2) compared to the neighboring eastern slope; and (3) to identify environmental parameters driving the observed differences.

## 2. Methods

### 2.1 DeSoto Canyon characteristics

The DeSoto Canyon cuts into the northwest Florida shelf and slope, ranging in depth from 400 – 3200 m (Fig 1). It is thought to be an inactive Canyon (Uchupi & Emery 1968, Bouma 1972) and is noted as a transition zone in seafloor sediment type (Antoine & Bryant 1968). Bottom substrate around the canyon to the west and east differs in size class and composition. To the west, sedimentation is dominated by siliclastic input from the Mississippi River. Bottom sediment primarily consists of quartz on the shelf, forming part of the Mississippi-Alabama-Florida Sand Sheet (Gould & Stewart 1955, Doyle & Sparks 1980). Continental slope sediments are rich in siliclastic clays and silts, in contrast to pelagic carbonate oozes that make up a majority of the deeper regions (Gould & Stewart 1955, Doyle & Sparks 1980, Balsam & Beeson 2003). East of the canyon, biogenic carbonate production highly influences sedimentation and forms the West Florida Sand Sheet on the mid-outer shelf, scaling down-slope to finer-grained West Florida Lime Mud (Doyle & Sparks 1980). Sediment accumulation rates range from ∼17 cm/ky (Emiliani et al. 1975) in the northwest area of the canyon to ∼10 cm/ky in the southeast (Emiliani et al. 1975, Nürnberg et al. 2008). When compensating for down-core compaction, accumulation rates reach 0.05 g/cm^2^/yr at ∼1850 m (Yeager et al. 2004). Particulate organic carbon (POC) can reach ∼0.67 - 1.67% of the top 18.5 cm of a core at depths of ∼1850 m (Yeager et al. 2004, Morse & Beazley 2008).

**Figure 1.**
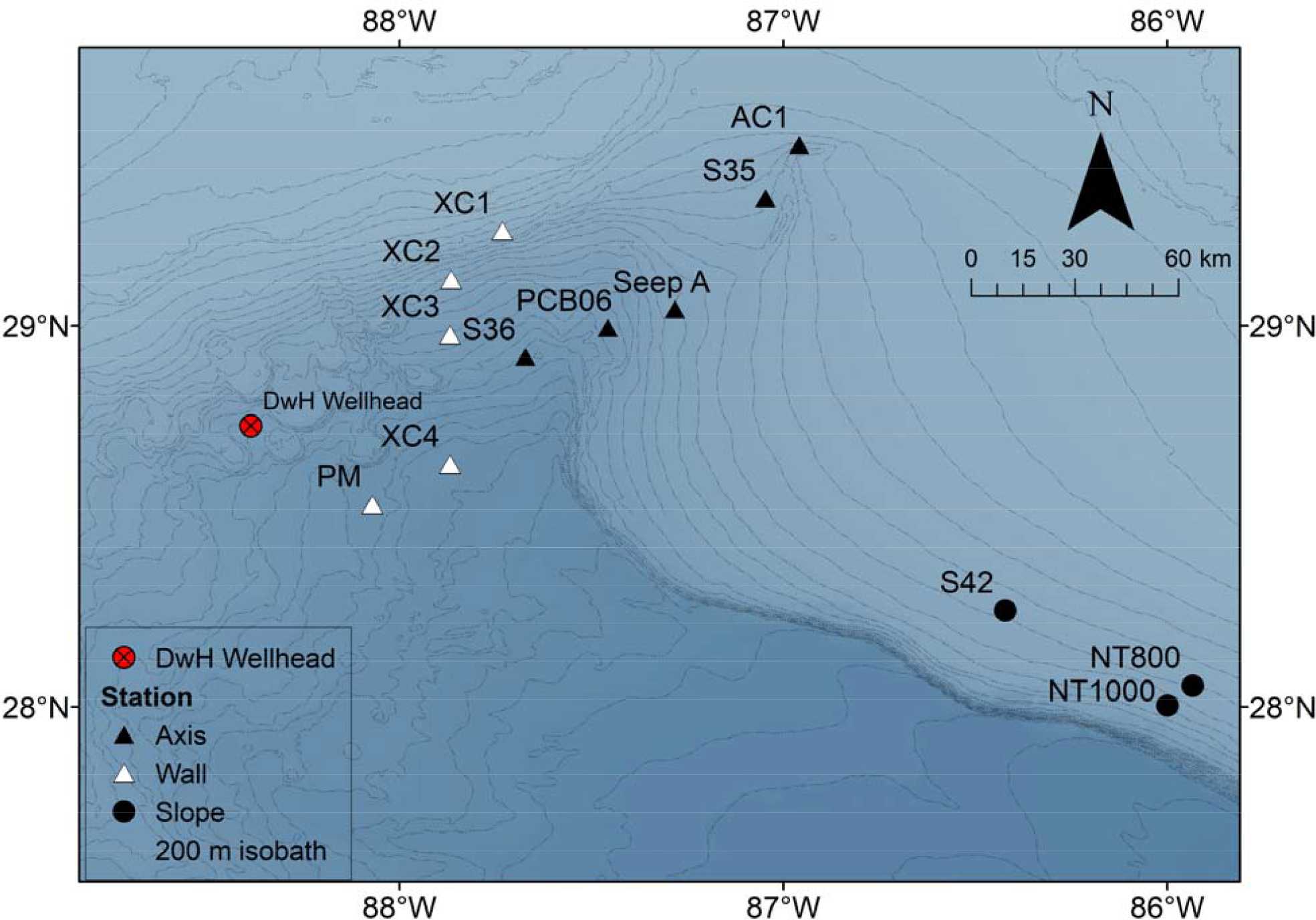
Bathymetric map of DeSoto Canyon sites sampled in 2014 relative to the position of the DwH wellhead. Ten sites traverse along the axis of the canyon (black triangles) and along the canyon wall (open triangles) and three are located on the adjacent slope (circles). Contour line depths are in meters.

### 2.2 Biological sample collection and processing

Sampling of ten sites within DeSoto Canyon was conducted as a part of the Gulf of Mexico Research Initiative (GOMRI) Deep-C Consortium during the May/June 2014 cruise aboard the *R/V Weatherbird II* cruise #WB1411 (Table 1, Figure 1). A comparable depth range of sampling sites was also targeted for the adjacent slope, but actual sampling was constrained by the compromises of a multi-PI cruise; thus, it was only possible to sample 3 non-canyon sites to the east of DeSoto Canyon. Sites within the DeSoto Canyon listed in Table 1 were selected to characterize spatial variability in canyon geomorphology, biogeochemistry, water column chemistry, and benthic communities along the canyon axis and canyon wall, following the 2010 Deepwater Horizon oil spill (Coleman et al. 2014), and to include DeSoto Canyon sites S35 and S36 of Wei et al (2010). Non-canyon sites were selected to compare the ecological and biogeochemical properties of the canyon with the open slope at the same depths and also to act as a control site outside the potential benthic footprint of the 2010 Deepwater Horizon (DwH) oil spill (Garcia-Pineda et al. 2013, Chanton et al. 2014), and to include one of the slope sites east of DeSoto canyon in Wei et al (2010), S42, that was most similar to S35 and S36 in that study.

**Table 1.** Station list with summary data for multicore deployments on the *R/V Weatherbird II*, in the DeSoto Canyon and adjacent slope in 2014. Three replicate deployments of the multicore were made at each station.

| Station | Lat (N) | Long (W) | Date (dd-mm-2014) | Depth (m) | Treatment |
| --- | --- | --- | --- | --- | --- |
| S35 | 29.3337 | -87.0502 | 1-Jun | 669 | Axis |
| PCB06 | 29.1950 | -87.4383 | 2-Jun | 1167 | Axis |
| S36 | 28.9163 | -87.6692 | 3-Jun | 1834 | Axis |
| Seep A | 29.0430 | -87.2825 | 7-Jun | 1114 | Axis |
| AC1 | 29.4745 | -86.9587 | 4-Jun | 464 | Axis |
| XC1 | 29.2482 | -87.7318 | 2-Jun | 485 | Wall |
| XC2 | 29.1210 | -87.8655 | 3-Jun | 1137 | Wall |
| XC3 | 28.9762 | -87.8683 | 4-Jun | 1510 | Wall |
| Peanut Mound (PM) | 28.5497 | -88.0862 | 9-Jun | 2045 | Wall |
| XC4 | 28.6365 | -87.8685 | 9-Jun | 2290 | Wall |
| NT800 | 28.0560 | -85.9335 | 30-May | 808 | Slope |
| NT1000 | 28.0040 | -85.9990 | 31-May | 978 | Slope |
| S42 | 28.2528 | -86.4217 | 31-May | 771 | Slope |

Three replicate deployments of an MC-800 Multicorer were conducted at each site. Each core had a diameter of 10 cm. Four cores from each deployment were sectioned on deck into 0-1, 1-5, 5-10 cm fractions and preserved whole in 10% formalin. In the laboratory, preserved samples were sieved through 300μm mesh and then transferred to 70% ethanol. Macrofaunal organisms (sensu stricto) were sorted using a dissecting microscope and identified to the lowest taxon possible, usually to class or order, and the dominant groups of bivalves, amphipods, cumaceans, and polychaetes were identified to the family level. Family level identification is considered sufficient to discern multivariate patterns in deep-sea ecosystems (Warwick 1988, Somerfield & Clarke 1995, Gesteira et al. 2003). Meiofauna that were >300μm (e.g., nematodes and harpacticoid copepods) were also identified and enumerated but left at the phylum to class level and excluded from analysis.

### 2.3 Sediment, water mass, and canyon terrain parameter measurement

Water column properties including temperature, salinity, oxygen, fluorescence (chlorophyll and colored dissolved organic matter (CDOM)), and turbidity were measured using the conductivity-temperature-depth (CTD) rosette aboard the *R/V Weatherbird II* at standard depths every 0.25 seconds after deployment from the surface until ∼ 10 m off the seafloor.

Bottom water conditions were obtained by averaging the parameters within ten meters of the bottom. Ocean-color data from (pixel size = ∼ 1 km^2^) were extracted from Visible Infrared Imaging Radiometer Suite (VIIRS) 8-day averages spanning mid-May to early June. Average surface chlorophyll concentration (SSC), photosynthetic aperture radar (PAR), and sea surface temperature (SST) were used as inputs to approximate depth-integrated net primary production (NPP) using a Vertical General Production Model (VGPM) (Behrenfeld & Falkowski 1997). Particulate organic carbon (POC) flux was approximated from NPP employing the exponential decay model of Lutz et al. (2007). More detailed methods on the ocean color data for the Gulf of Mexico are provided in Biggs et al. (2008).

Sediment parameters of total organic carbon (TOC), total organic nitrogen (TON), and grain size were measured from the 0-5 cm depth section of a spare core from each deployment using a 30-cc syringe. Carbon and nitrogen samples were treated with 10% HCl to remove carbonates. Subsequently, samples were freeze dried, ground, and sealed in tin cups for combustion in a ThermoQuest CE Instrument NC2500 Analyzer. Percent carbon and nitrogen were measured on a Thermo Fischer Scientific Delta Plus XP Isotope Ratio Mass Spectrometer. Grain size subsamples of the same sediment core were taken from 0-5 cm and measured whole for granulometry. The samples were dried to in an oven at 100°C overnight, ground to a powder, and then treated with 15 ml 30% H_2_O_2_ and 15 ml 10% HCl to remove organic matter and carbonates respectively (Jackson 1969). The powdered sediment was then suspended in water and the grain size distribution was measured via laser diffraction using a Mastersizer 2000MU Hydro. The samples were characterized by their percent clay (<8μm), silt (8-63 μm), and sand (>63 μm) volume proportions, defined after Konert and Vandenberghe (1997). Characteristics of the canyon sediment surface including slope, aspect, and rugosity (surface roughness) were dervied from the bathymetry layer using the Benthic Terrain Modeler Tool (Rinehart et al. 2004) in ArcMap 10.6.1. Slope was calculated in degrees using the 3 x 3 cell window (Burrough et al. 2015). Aspect calculates the downslope direction, measured clockwise in degrees from 0 (north) to 360 (north) of each cell in relationship to its neighbors. It is derived from the z (bathymetry) values in a 3 x 3 cell window (Burrough et al. 2015). As a circular variable, it was converted into two parameters, northness (computed as cos(aspect)) and eastness (computed as sin(aspect)). These parameters characterize sites that took a north-south aspect and sites of an east-west aspect. A full list of environmental variables with data ranges can be found in Table 2.

**Table 2.** Summary of environmental factors sampled in the DeSoto Canyon with ranges of values for each parameter across all samples.

| Variable type | Name | Source | Sample Availability |  | Range | Unit |
| --- | --- | --- | --- | --- | --- | --- |
|  |  |  | Canyon | Slope |  |  |
| Seafloor/Terrain | Slope | Bathymetry | X |  | 0.16 – 4.33 | degrees |
|  | Aspect - northness | Bathymetry | X |  | -1.0 – 1.0 | degrees |
|  | Aspect-eastness | Bathymetry | X |  | -1.0 – 1.0 | degrees |
| Sediment environment | %carbon | Core sub-sample | X | X | 1.30 – 2.67 | % |
|  | %nitrogen | Core sub-sample | X | X | 0.12 – 0.36 | % |
|  | %sand | Core sub-sample | X | X | 14.81 – 65.04 | % |
|  | %silt | Core sub-sample | X | X | 30.43 – 74.93 | % |
|  | %clay | Core sub-sample | X | X | 0.14 – 29.50 | % |
| Water mass | Salinity | CTD rosette | X | X | 34.89 – 35.27 | PSU |
|  | Temperature | CTD rosette | X | X | 4.27 – 10.61 | deg C |
|  | O<sub>2</sub> sat | CTD rosette | X | X | 3.81 – 6.72 | mg/l |
|  | Fluorometry Eco-Afl and CDOM | CTD rosette | X | X | 2.95 – 9.04 | mg/m<sup>3</sup> |
|  | Turbidity | CTD rosette | X | X | 0.18 – 8.08 | NTU |
|  | Particulate Organic Carbon | Remote sensing | X | X | 7.90 – 34.35 | mg C/m<sup>2</sup>/day |

### 2.4 Statistical comparisons

For all statistical analyses, the four cores from each deployment were combined as one sample, with the deployments as the replicates for that site. The sampling constraints described above resulted in the range of depths of the non-canyon sites being only a subset of the depths sampled within the canyons. This prevented a balanced design for a comparison of within canyon vs. non-canyon sites and so data were analyzed in two phases. Since depth (and its correlates) is known to be a strong structuring factor in the deep sea (reviewed in Rex and Etter 2010), in phase I all the sampling stations within the canyon (depth range 464 – 2290 m) were analyzed, to determine the depth structuring of the canyon communities. Then, to avoid the confounding of depth, in phase II all samples in the depth group determined in phase I that overlapped with the sampled depth range of the non-canyon sites (771-978 m), were used in the comparisons among canyon wall, canyon axis, and adjacent non-canyon slope habitat types.

Differences in macrofaunal community abundances and diversity metrics were tested as a product of the following fixed factors in a one-way design: (1) *a posteriori* canyon depth groups (464 – 485 m vs 669 – 1834 m vs > 2000 m) and (2) habitat type (canyon wall vs canyon axis vs slope). Due to the large differences in sample size among depth and habitat groups, non-parametric Kruskal-Wallis tests were conducted to test for differences, with Bonferoni-adjusted Dunn’s pairwise post-hoc analysis.

For multivariate analysis, community structure was depicted via cluster analysis and non-metric multidimensional scaling (NMDS). ANOSIM, based on Bray-Curtis similarity, was used to test the *a priori* habitat types in phase II. Due to the imbalance in sample size between habitat types (canyon axis and slope sites outnumber the canyon wall sites in this depth range), biases may be encountered in the ANOSIM (Anderson & Walsh 2013), thus samples were removed from the largest groups at random to match the smallest group. To ascertain which taxa were driving observed differences between communities in each habitat type, a similarity percentage (SIMPER) analysis was employed. Distance-based linear modeling (DISTLM) (Anderson et al. 2008), was used to find the optimal combination of abiotic factors that significantly correlated with community structure. Prior to DISTLM analyses, environmental variables were normalized and plotted pairwise using draftsmen plots. Log-transformations were applied to highly skewed individual variables and highly collinear factors (>90%) removed. Interpolation of the environmental parameters was conducted to replace missing replicates and to run analyses.

Environmental variables were first analyzed individually (marginal tests) and then the *BEST* selection procedure was employed to select the optimal model based on the small sample adjusted Akaike Information Criterion (AICc) for all possible combinations of environmental predictor variables. AICc was employed because it was formulated to deal with situations where the number of observations (N) to the number of variables (v) is < 40 (Burnham & Anderson 2004) as in the case of this dataset (N = 39, v ≤ 13, N/v = 3.0).

The environmental data used for the input into the DISTLM was not the same for both phases of analyses. The DISTLM for the phase I within canyon analyses included all sediment, water mass, and terrain parameters. However, slope and terrain were unavailable for the non-canyon slope sites because high-resolution bathymetry was not available (Table 2), so terrain parameters were not included for the canyon axis vs. wall vs. slope DISTLM in phase II.

For all tests, differences at p < 0.05 were considered significant. All statistical comparisons were conducted in R (R Core Team, 2019) and multivariate analyses were conducted using in PRIMER v 7.0.13 (Clarke & Gorley 2015).

## 3. Results

### 3.1 Macrofaunal abundance and diversity within the DeSoto canyon

Within the DeSoto Canyon, a total of 6637 individuals were identified to the lowest taxonomic level possible, most often family. Polychaetes (49.09 – 77.84%) were the most abundant taxonomic group, followed by tanaids (2.27 – 16.46%), bivalves (2.84 – 13.53%), nemertean worms (2.32 – 5.80%), and amphipods (0.32 – 4.88%) (Table 3). Groups with otherwise low individual proportions, when aggregated to the phylum and subphylum level, exhibit large relative abundance. These include other molluscs (scaphopods, gastropods, and cavoliniids), which contained proportions 2.37 – 16.99%, and other crustaceans (isopods and cumaceans), which had a relative abundance of 0.23 – 17.99%. Relative contribution from each taxonomic group changed by site (Table 3). Anomalously high abundances compared to the mean were found for bivalves at XC3, other molluscs at XC2, and for tanaids at PM and S35. S36, PM and XC4 had high values for other crustaceans compared to the other sites.

**Table 3.**
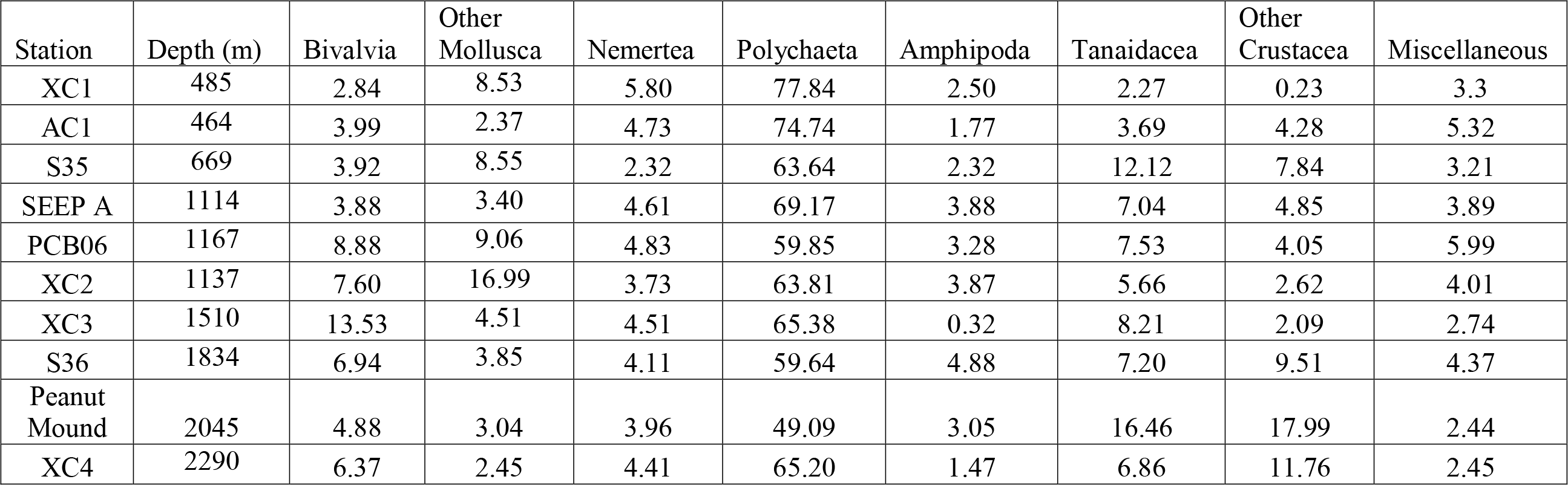
Major taxonomic group proportions for Desoto Canyon macrofauna by station.

By depth, the highest average abundance was observed at 485 m with a continual decrease throughout the canyon (Fig 2A). A significant relationship was found with depth (p = 0.003). Mean richness formed a significant (p = 0.0006) parabolic relationship with depth reaching a maximum around 1100 m (Fig 2B). Average Pielou’s evenness increased with depth, ranging between 0.75 – 0.90 (Fig 2C) and was also found to have a significant increase with depth (p = 0.044).

**Figure 2.**
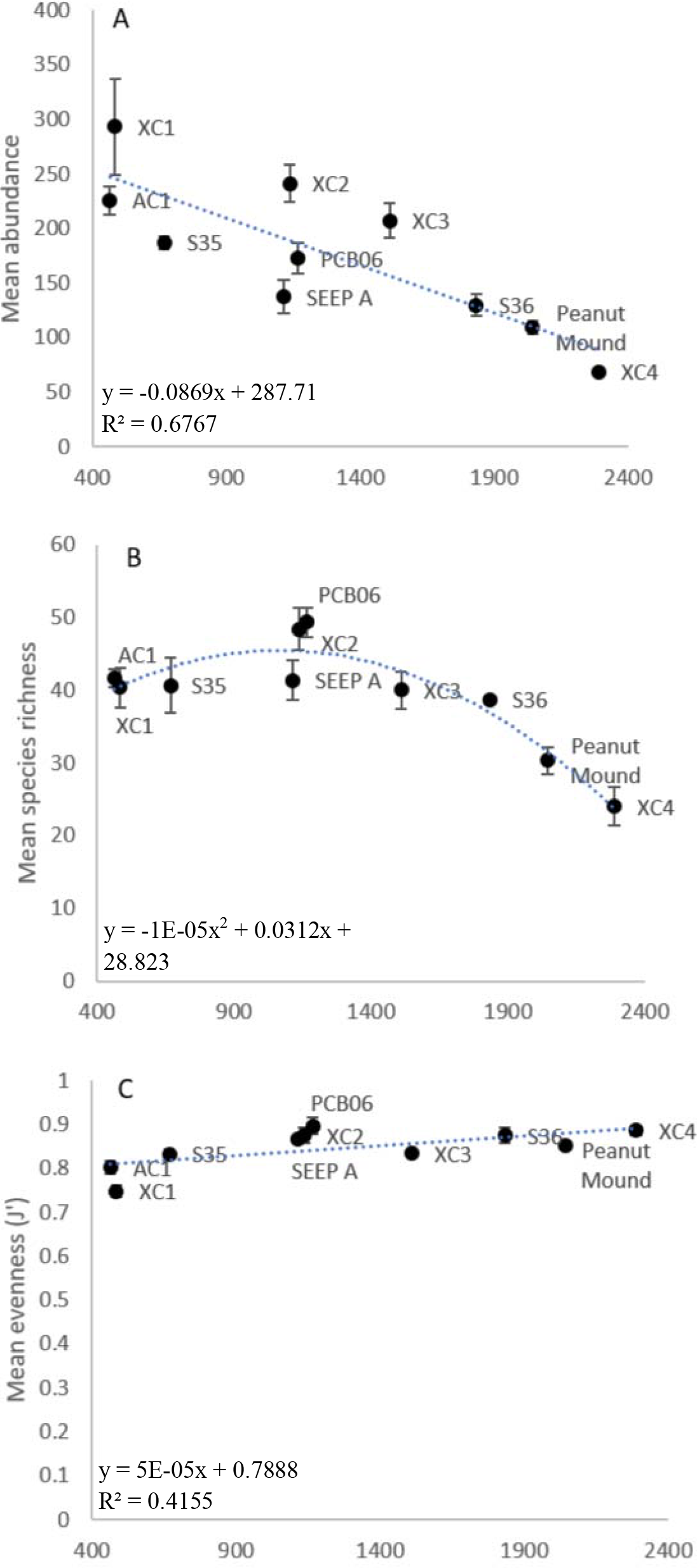
Mean abundance and diversity metrics within the DeSoto Canyon ordered by depth. A) Abundance (F_(1, 8)_ = 16.75). B) Species richness (F_(2, 7)_ = 25.56). C) Pielou’s evenness (F_(1, 8)_ = 5.686). Error bars are standard error of the mean.

### 3.2 Community structure within the DeSoto Canyon

Three depth assemblages were identified through the cluster analysis of the within canyon macrofauna (Fig 3A) and depicted via non-metric multidimensional scaling (Fig 3B): Assemblage Group I included the shallowest canyon sites (464 – 485 m), Group II included the bulk of the canyon sites (670 – 1834 m), and Assemblage Group III included the deepest sites (> 2000 m). Among these *a posteriori* depth groups, all main effect tests of abundance and within canyon diversity metrics showed significant differences overall (Fig. 4). Abundance was significantly different among all pairs of depth groups (Fig 4A). Pairwise comparison of depth groups for species richness only found differences for the > 2000 m sites, which had lower richness compared to either of the other two depth groups (Fig 4B). Evenness was lower for the 464 – 485 m sites compared to the deeper depth groups (Fig 4C).

**Figure 3.**
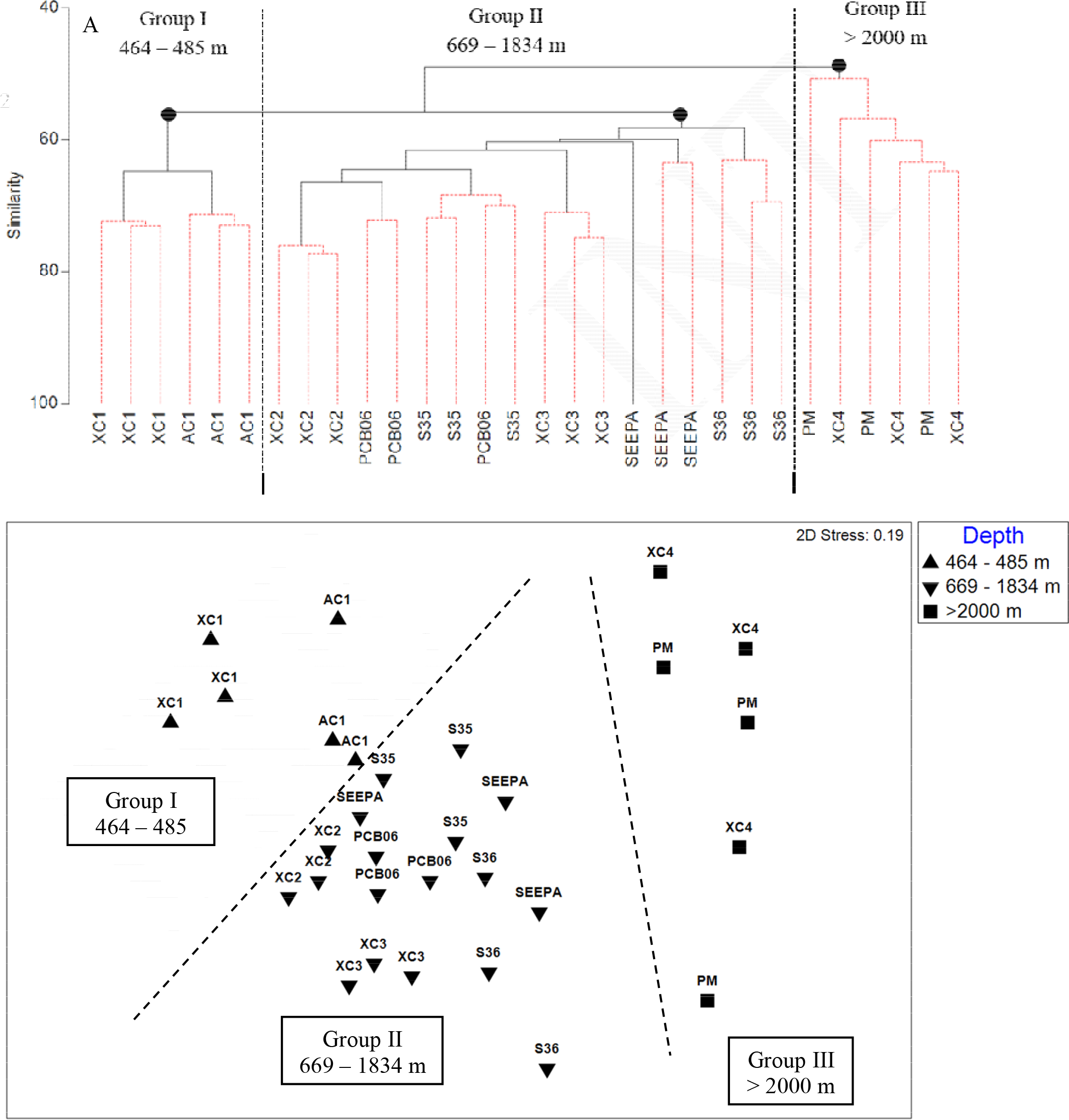
A) Cluster analysis based on root-transformed abundances of DeSoto Canyon macrofauna. The black dots indicate nodes of significant clusters (Pairwise ANOSIM R = 0.526 – 0.904, p 0.002). B) Non-metric multidimensional scaling of DeSoto Canyon macrofauna.

**Figure 4.**
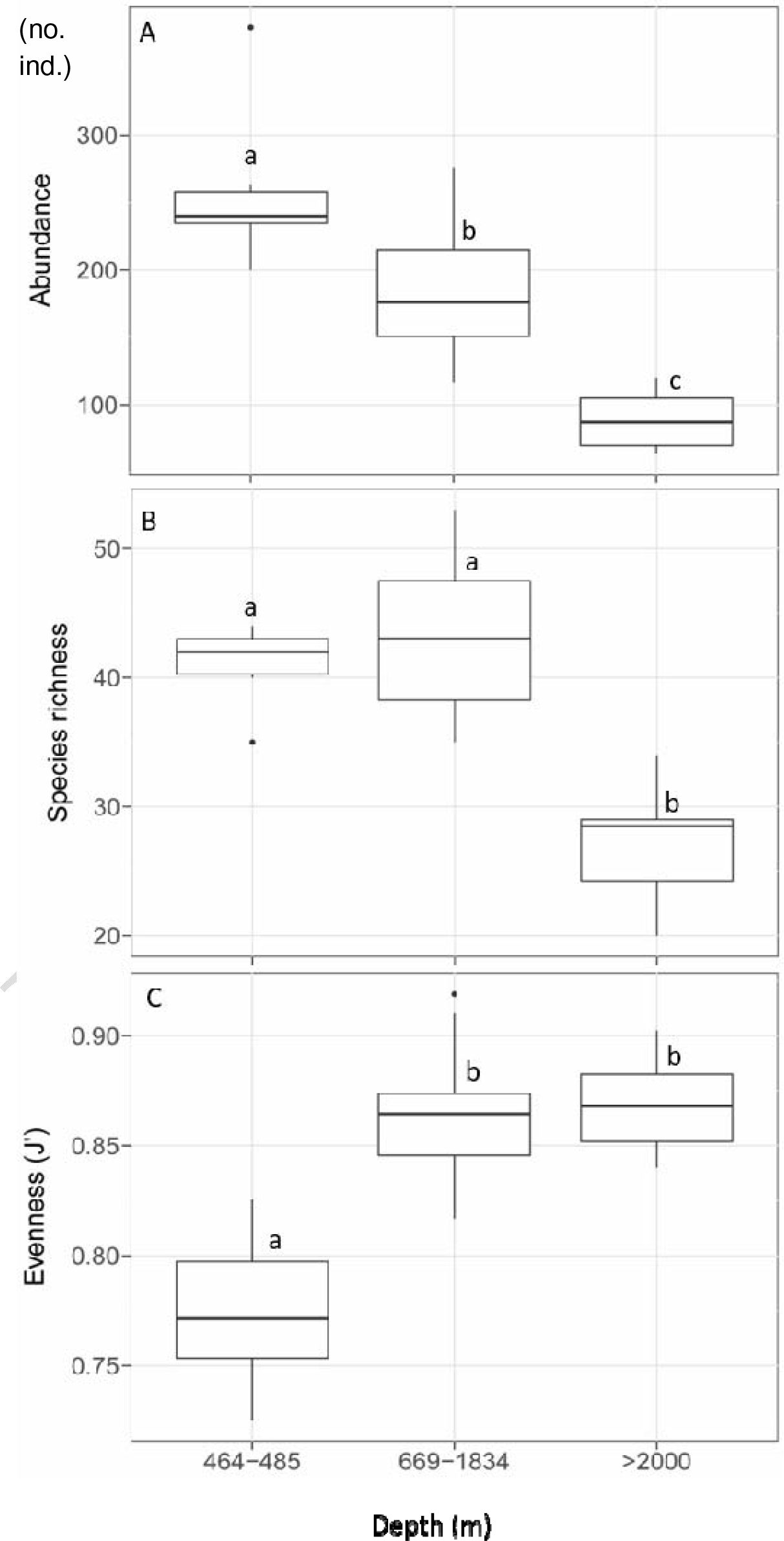
Diversity metric comparisons among *a posteriori* depth groups for DeSoto Canyon macrofauna (A) abundance (χ^2^ = 19.148; p < 0.001), (B) species richness (χ^2^ = 14.28; p < 0.001), and (C) Pielou’s evenness (χ^2^= 13.278; p = 0.0013). Shared letter indicates no statistical difference between depth groups(P>0.05)

Only those environmental variables with low collinearity with other variables (R^2^ < 0.90) were included in the DISTLM. Temperature had a high correlation with POC and oxygen saturation, so it was removed prior to analysis. Of the remaining 14 variables available for the DISTLM for communities within the canyon, 10 were found to be significant as indicated by the marginal tests (Table 4). AICc values computed for top models spanned a narrow range (204.09 – 204.97) suggesting rather equivalent models explained the variation in community structure, as typically a difference of 2 units between models indicates separate models (Burnham & Anderson 2004, Anderson et al. 2008). The top model selected by DISTLM was a combination of oxygen saturation and fluorometry (R^2^ = 0.2556). The top models all included fluorometry, and fluorometry by itself received an AICc value only 0.8 less than the best model. The top five models contained some combination of oxygen, salinity, and/or percent organic carbon, with fluorometry. For sediment and terrain parameters, percent organic carbon and slope were the only to appear in the top models, with relatively similar fits, R^2^ = 0.3085 and 0.3014 respectively.

**Table 4.** DISTLM marginal tests and overall best solutions for the environmental factors compared to macrofaunal assemblage structure within the DeSoto Canyon.

| Variable No. | Variable | SS(trace) | Pseudo-F | P | Prop. |
| --- | --- | --- | --- | --- | --- |
| 1 | Salinity | 3227.2 | 3.5318 | **0.002** | 0.11201 |
| 2 | O<sub>2</sub> saturation [mg/l] | 4141.9 | 4.701 | **0.001** | 0.14376 |
| 3 | Fluorometry Eco-afl<br>mg/m<sup>3</sup> | 4887.9 | 5.7207 | **0.001** | 0.16965 |
| 4 | Fluorometry CDOM<br>mg/m<sup>3</sup> | 1590.3 | 1.6358 | 0.069 | 0.055197 |
| 5 | POC | 4593.4 | 5.3106 | **0.001** | 0.15943 |
| 6 | Turbidity | 2916.2 | 3.1531 | **0.002** | 0.10121 |
| 7 | %carbon | 4295.8 | 4.9062 | **0.001** | 0.1491 |
| 8 | %nitrogen | 3175.6 | 3.4684 | **0.001** | 0.11022 |
| 9 | %sand | 2152.3 | 2.2606 | **0.009** | 0.074703 |
| 10 | %silt | 1698.3 | 1.7538 | **0.044** | 0.058944 |
| 11 | %clay | 1226.8 | 1.2452 | 0.17 | 0.042578 |
| 12 | Aspect-northness | 800.71 | 0.80038 | 0.691 | 0.027791 |
| 13 | Aspect-eastness | 1400.6 | 1.4306 | 0.118 | 0.048611 |
| 14 | Slope | 2707 | 2.9035 | **0.001** | 0.093954 |
| Overall Best Solutions |  |  |  |  |  |
| AICc | R<sup>2</sup> | RSS | No. Variables | Selections |  |
| 204.09 | 0.25555 | 21449 | 2 | 2,3 |  |
| 204.25 | 0.31549 | 19722 | 3 | 1-3 |  |
| 204.4 | 0.24777 | 21673 | 2 | 1,3 |  |
| 204.55 | 0.3085 | 19923 | 3 | 2,3,7 |  |
| 204.62 | 0.24227 | 21832 | 2 | 3,7 |  |
| 204.71 | 0.30498 | 20025 | 3 | 2-4 |  |
| 204.86 | 0.30142 | 20127 | 3 | 2,3,14 |  |
| 204.89 | 0.16965 | 23924 | 1 | 3 |  |
| 204.91 | 0.30025 | 20161 | 3 | 2,3,5 |  |
| 204.97 | 0.23345 | 22086 | 2 | 3,5 |  |

The top model is plotted in the dbRDA plot (Fig 5). Deeper sites (Group III) and most of the mid-slope sites (Group II) tended to fall higher along dbRDA axis 2. The shallowest sites and S35 were differentiated along both axes. The dbRDA1 axis, explained 66.6% of the fitted variation, but 17% of the community structure variation. Fluorometry had the strongest relationship (0.864) with the first axis. The dbRDA2 axis, accounting for 33.4% of the fitted and 8.5% of the overall variation, had the strongest association with oxygen saturation (0.864).

**Figure 5.**
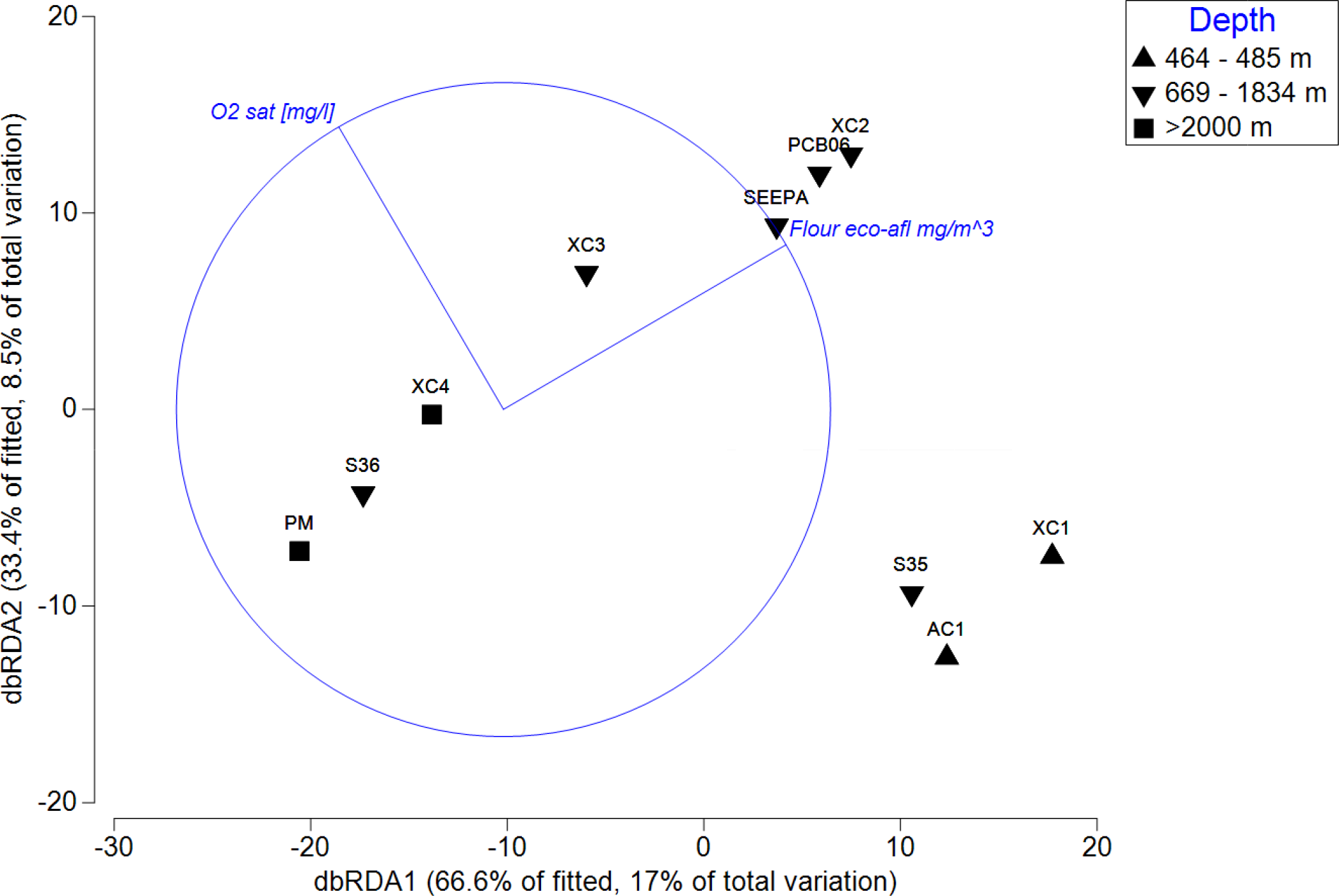
Distance-based redundancy analysis (dbRDA) plot of the top DISTLM model of community structure and environmental variables within DeSoto Canyon.

### 3.3 DeSoto Canyon axis and wall vs. non-canyon slope: macrofaunal variation and abiotic factors

Macrofaunal proportions by total individuals of major groups were reasonably comparable across habitats in the canyon and on the adjacent slope (Fig 6). Polychaetes dominated with proportions ranging from 58.41 – 64.54%, with slightly more in the canyon habitats (63 - 65%) than the slope (58%). The next most abundant groups varied depending on habitat and were generally the tanaids (8.84 – 8.72%) and bivalves (5.52 – 10.33%). Tanaids held relatively similar proportions between habitats while bivalves exhibited higher proportions on the canyon wall compared to the canyon axis and adjacent slope. Remaining groups held proportions approximately 6% or less though macrofauna in too low of abundance to form their own group, termed ‘other’, exhibited a combined proportion of 9.17% on the continental slope.

**Figure 6.**
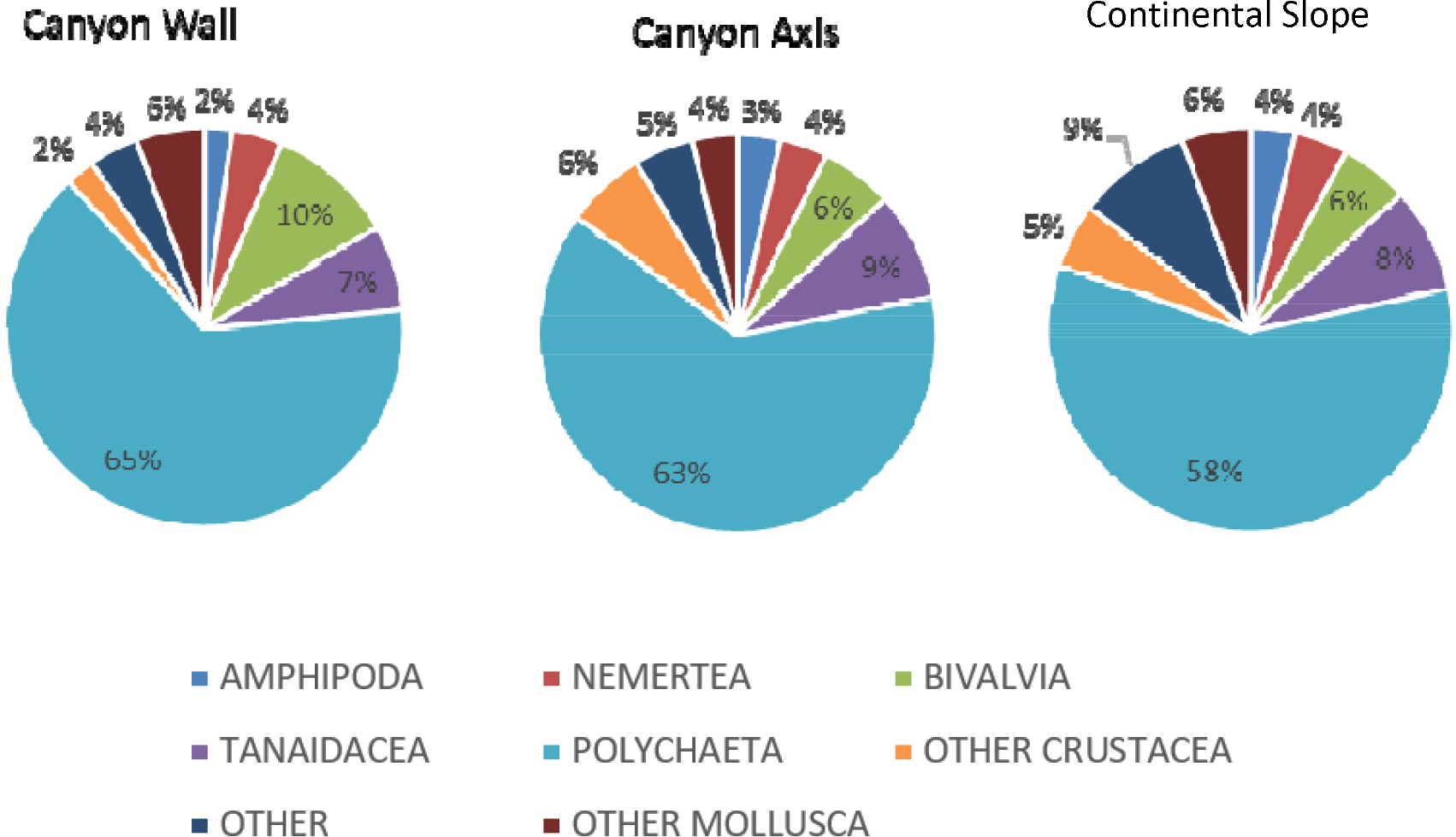
Relative abundance of major taxonomic groups by overall totals in the DeSoto Canyon habitats (wall, axis) compared to the adjacent continental slope.

Global tests of abundance and diversity metrics of the three habitat types only detected differences for abundance (p < 0.001). Abundance was highest on the canyon wall, followed by the axis, and then the slope (Fig 7A). Pairwise comparisons of the canyon axis, wall and the adjacent slope were all significantly different (p < 0.05). No differences were found among habitats for species richness (Fig 7B) nor evenness (Fig 7C).

**Figure 7.**
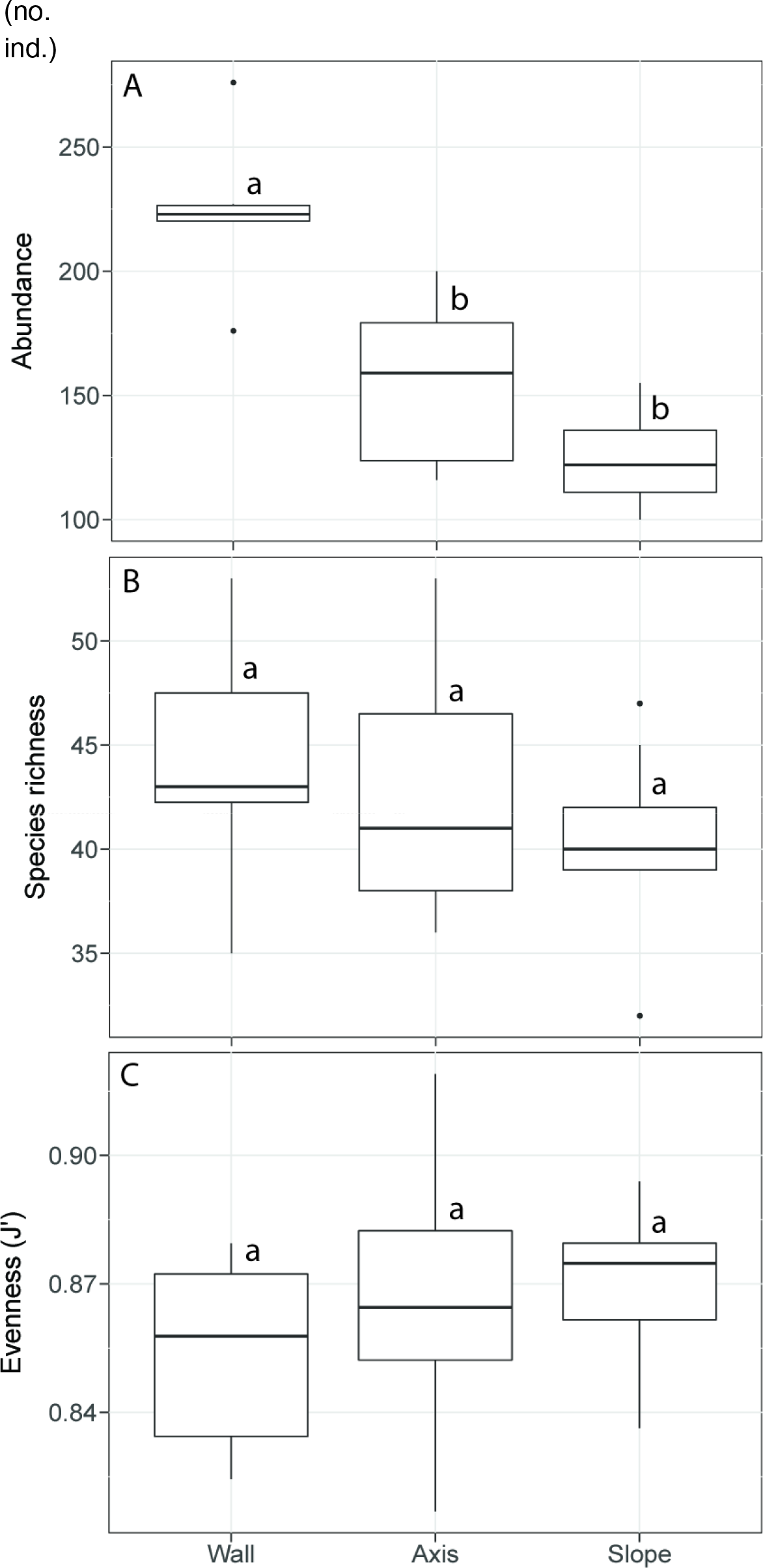
Diversity metrics comparing canyon habitat Group II axis and wall sites (669 – 1510 m) and adjacent slope (771 – 978 m). A) Abundance (χ^2^= 15.72; p < 0.001). B) Species richness (χ^2^ = 1.3324; p = 0.5137). C) Pielou’s evenness (χ^2^ between depth groups (p > 0.05). = 1.4951; p = 0.4735). Shared letter indicates no statistical difference between depth groups (p > 0.05).

Of the 12 parameters available for comparison between habitat types, all but chlorophyll-based fluorescence, POC, and percent silt showed significant differences among the habitat types. Temperature was lower in the canyon compared to the adjacent slope (Fig 8A). Salinity was significantly higher in the canyon compared to the slope (Figure 8B). Oxygen saturation was significantly higher on the canyon wall (6.05 – 6.63 mg/l) and canyon axis (4.21 – 6.69 mg/l), compared to the slope (4.79 – 5.56 mg/l) (Fig 8C). CDOM fluorescence was higher on the canyon wall than the slope (Fig 8E). Turbidity was higher in the canyon (Fig 8F). Organic matter was significantly lower in the canyon for sediment percent carbon and percent nitrogen (Fig 8H-I). Sediment percent sand of the canyon axis was significantly higher than the slope (Fig 8J). Percent clay in the canyon wall and slope sites were higher than the canyon axis (Fig 8L).

**Figure 8.**
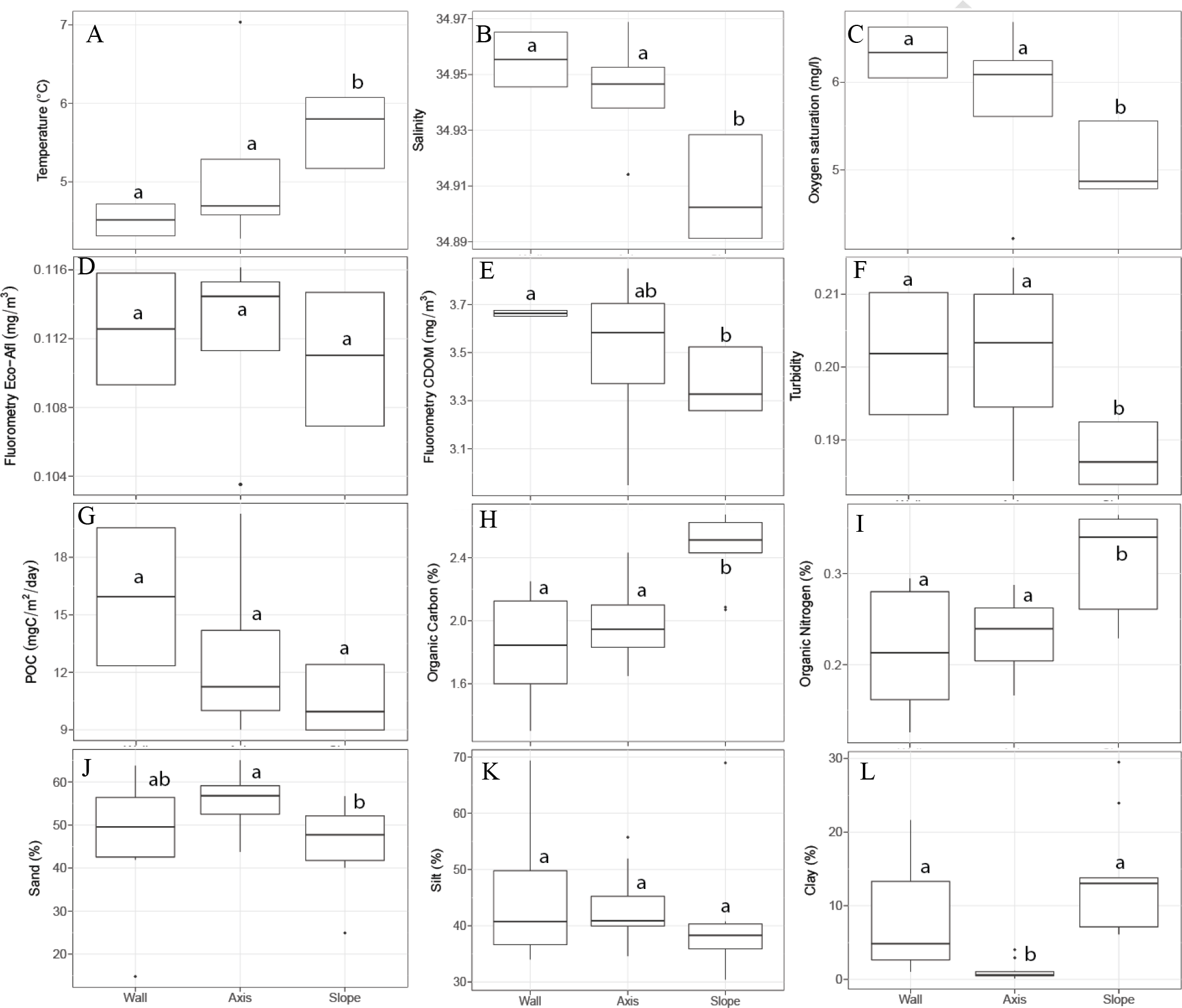
Boxplots of environmental factors across habitat types (canyon wall, axis and adjacent slope).Shared letter indicates no statistical difference between depth groups (p > 0.05). A) Temperature (χ^2^ =8.125, p = 0.01721). B) Salinity (χ^2^ = 13.903, p < 0.001). C) Oxygen saturation (χ^2^= 8.125, p = 0.01721). D) Fluorometry Eco-Afl (χ^2^= 1.95, p = 0.3772). E) Fluorometry CDOM (χ^2^= 7.1861, p = 0.02751). F) Turbidity (χ^2^ = 10.761, p = 0.004605). G) POC flux (χ^2^ = 5.7778, p = 0.05564). H) Organic carbon (χ^2^ = 12.568, p = 0.001866). I) Organic nitrogen (χ^2^ = 8.3891, p = 0.01508). J) %sand (χ^2^ = 6.9524, p = 0.03092). K) %silt (χ^2^ = 2.7557, p = 0.2521). L) %clay (χ^2^ = 18.015, p < 0.001).

### 3.4 Community structure and relation to environmental variables of canyon and non-canyon habitats

A one-way ANOSIM was significant for community structure across habitats (p < 0.001, Table 5). All pairwise comparisons of habitat types were also significant, indicating differences between all three habitats (Table 5). Community structure differences of canyon axis and wall and slope sites, depicted via NMDS in Figure 9, also portray a west-to-east longitudinal gradient moving from left to right across the ordination. Between canyon habitats, SIMPER analysis results revealed an average dissimilarity of 36.5% (Table S1). Taxa contributing the most to differences (> 2%) included clams of the family Thyasiridae, numerous deposit feeding groups spanning longosomatids, maldanids, syllids, paraonids, and cirratulid polychaetes, as well as aplacophorans. Carnivorous and omnivorous polychaetes of the Families Hesionidae and Sigalionidae were identified as well. Between slope and wall habitats, dissimilarity averaged 43.4% and many of the same groups differentiated community structure but also included polynoids, fauveliopsids, and capitellids (Table S1). Taxa differentiating canyon axis and adjacent slope habitats (average dissimilarity 39.0%) were fauveliopsid, syllid, sigalionid, maldanid, and paralacydoniid polychaetes. Additionally, malletiid bivalves and various cnidarians made contributions (Table S1).

**Figure 9.**
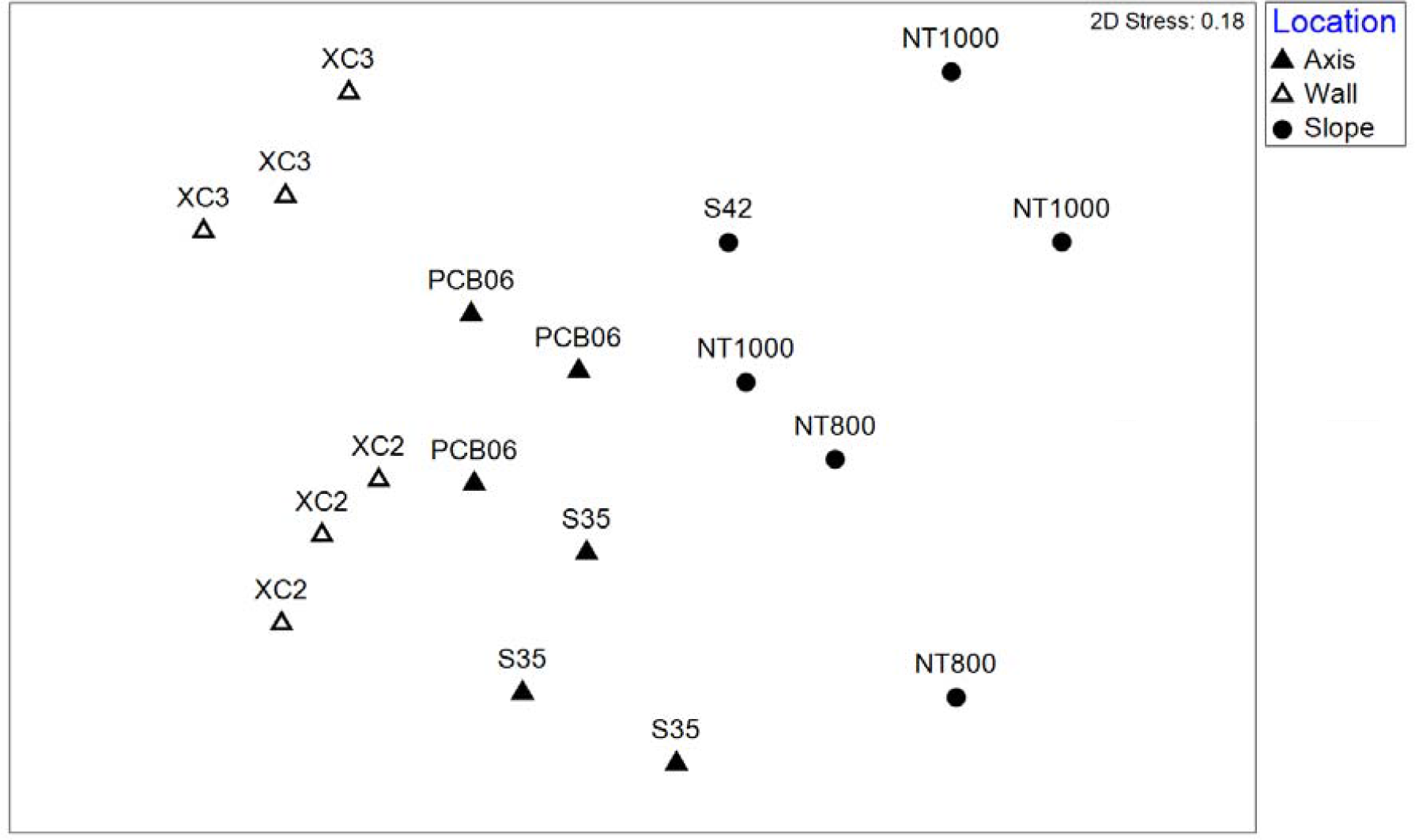
Non-metric multidimensional scaling of group II canyon axis and wall sites at depths of 669 – 1510 m compared to slope sites at 771 – 978 m.

**Table 5.** One-way ANOSIM with pairwise comparisons of community structure among and between habitat types in the canyon (wall and axis, 669 – 1834 m) and adjacent slope (771 – 978 m). Bolded values indicate significant differences between groups (p<0.05).

| Canyon Wall vs. Axis vs. Slope | $R^2$ | p-value | Permutations |
| --- | --- | --- | --- |
| Global test | 0.483 | **0.001** | 999 |
| Pairwise groups |  |  |  |
| Slope, Axis | 0.485 | **0.004** | 462 |
| Slope, Wall | 0.757 | **0.002** | 462 |
| Axis, Wall | 0.291 | **0.030** | 462 |

The environmental factors of temperature and salinity were removed from consideration prior to DISTLM to avoid model bias from high correlation with other variables, leaving 10 variables available for analysis of community structure differences between habitat types. Of these all were significantly correlated with macrofauna community structure except percent sand and percent silt (Table 6). AICc values spanned a small range (180.4 – 181.41). The BEST model selected by DISTLM to explain most of the macrofaunal community variation included only 2 factors, oxygen saturation and POC flux, explaining 20.7% of macrofaunal community variation (Table 6). Water mass parameters exclusively comprised the top 9 models that explained the most community variation and most of the models contained oxygen saturation. In fact, the fifth best model included oxygen by itself with an AICc value only 0.84 higher than the top model. The 10^th^ model was the only model to contain a sediment parameter, percent carbon, which was paired with POC flux. The first axis of the dbRDA plot of the top model (Fig 10) explained 64.2% of the fitted variation 13.3% of the total) and was strongly correlated with oxygen saturation (-0.803). The second axis accounted for 35.8% of the fitted variation and 7.4% overall and was most strongly correlated with POC flux (0.803).

**Table 6.**
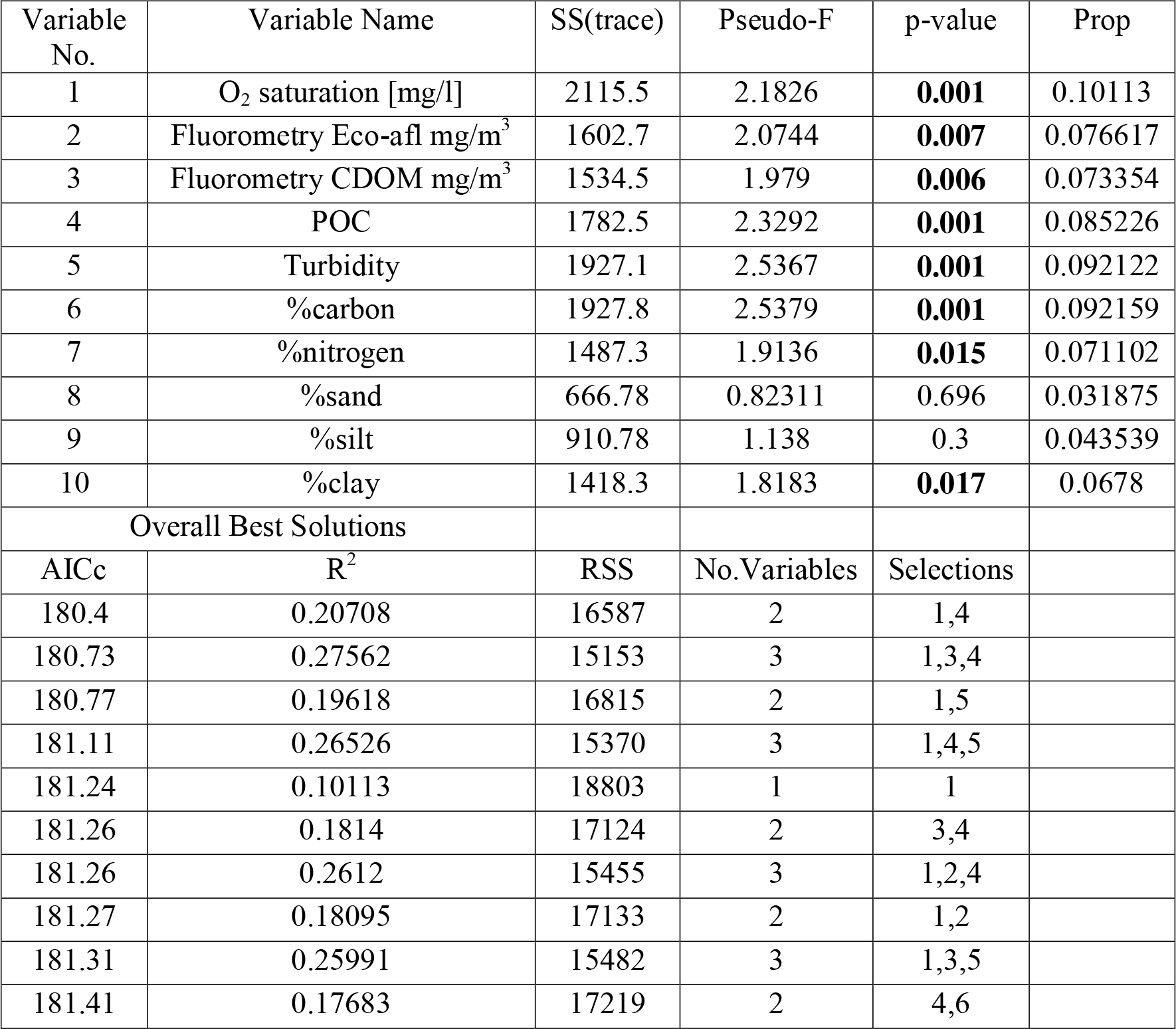
. DISTLM marginal tests and overall best solutions for the environmental factors compared to macrofaunal assemblage structure among the axis and wall canyon macrofauna communities at depths of 669 – 1834 m compared to the adjacent non-canyon slope at 771 – 978 m depth.

| Variable No. | Variable Name | SS(trace) | Pseudo-F | p-value | Prop |
| --- | --- | --- | --- | --- | --- |
| 1 | O<sub>2</sub> saturation [mg/l] | 2115.5 | 2.1826 | **0.001** | 0.10113 |
| 2 | Fluorometry Eco-afl mg/m<sup>3</sup> | 1602.7 | 2.0744 | **0.007** | 0.076617 |
| 3 | Fluorometry CDOM mg/m<sup>3</sup> | 1534.5 | 1.979 | **0.006** | 0.073354 |
| 4 | POC | 1782.5 | 2.3292 | **0.001** | 0.085226 |
| 5 | Turbidity | 1927.1 | 2.5367 | **0.001** | 0.092122 |
| 6 | %carbon | 1927.8 | 2.5379 | **0.001** | 0.092159 |
| 7 | %nitrogen | 1487.3 | 1.9136 | **0.015** | 0.071102 |
| 8 | %sand | 666.78 | 0.82311 | 0.696 | 0.031875 |
| 9 | %silt | 910.78 | 1.138 | 0.3 | 0.043539 |
| 10 | %clay | 1418.3 | 1.8183 | **0.017** | 0.0678 |
| Overall Best Solutions |  |  |  |  |  |
| AICc | R<sup>2</sup> | RSS | No. Variables | Selections |  |
| 180.4 | 0.20708 | 16587 | 2 | 1,4 |  |
| 180.73 | 0.27562 | 15153 | 3 | 1,3,4 |  |
| 180.77 | 0.19618 | 16815 | 2 | 1,5 |  |
| 181.11 | 0.26526 | 15370 | 3 | 1,4,5 |  |
| 181.24 | 0.10113 | 18803 | 1 | 1 |  |
| 181.26 | 0.1814 | 17124 | 2 | 3,4 |  |
| 181.26 | 0.2612 | 15455 | 3 | 1,2,4 |  |
| 181.27 | 0.18095 | 17133 | 2 | 1,2 |  |
| 181.31 | 0.25991 | 15482 | 3 | 1,3,5 |  |
| 181.41 | 0.17683 | 17219 | 2 | 4,6 |  |

**Figure 10.**
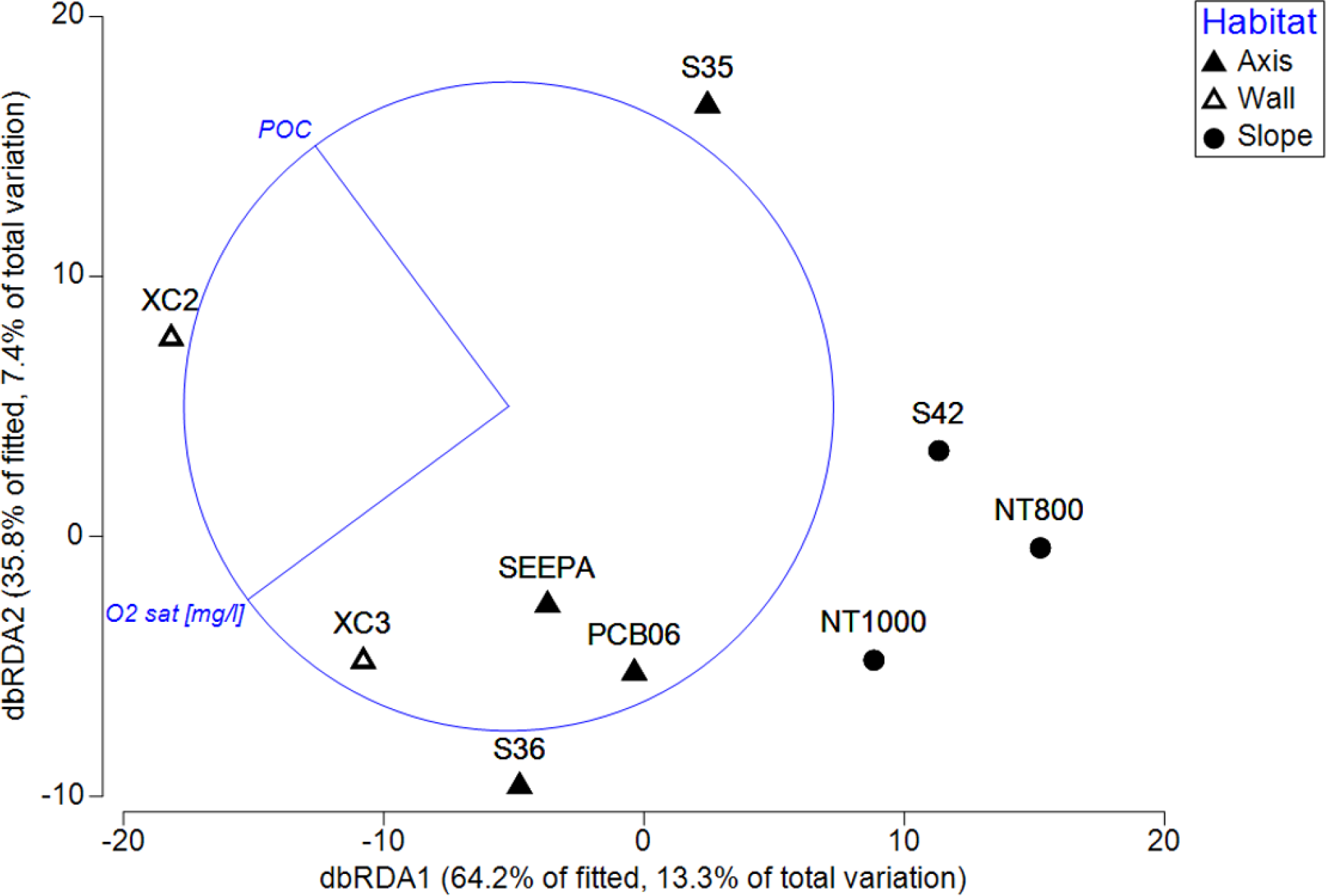
dbRDA plots of the canyon at 669 – 1834 m compared to the adjacent slope at 771 – 978 m.

## 4. Discussion

### 4.1 Influence of the DwH spill on the DeSoto canyon

The primary goal of this study was to examine spatial variability in macrofaunal communities within the DeSoto Canyon that may have been missed at the coarse sampling scales previously undertaken. However because this study was undertaken <4 years after the Deepwater Horizon (DwH) Oil Spill, we must first consider what effect, if any, the spill had on the DeSoto Canyon fauna. Starting in April 2010, the DwH spill release 130 M gal of crude oil and natural gas from a depth of 1500 m (McNutt et al. 2012). Of the total oil, 3.0-4.9% (1.6 to 2.6 x 10^10^ g) is estimated to have deposited to the deep seafloor in a 8400 km^2^ footprint, with the highest concentration found in a 3200 km^2^ area immediately around the wellhead (Chanton et al. 2014, Valentine et al. 2014). Small oil droplets and dissolved oil and gas formed plumes at two known depths, 50 – 500 m and 1,100 – 1500 m (Camilli et al. 2010, Socolofsky et al. 2011, Valentine et al. 2014). Where the plumes intersected the continental slope, hydrocarbons deposited.

Hydrocarbons at the surface and persisting in the water column structured microbial blooms (Hazen et al. 2010, Valentine et al. 2010, Kessler et al. 2011, Redmond & Valentine 2012, Mason et al. 2014a, Kleindienst et al. 2015) whose products aggregated with unprocessed hydrocarbons, bacterial products, and phytoplankton (Passow et al. 2012, Ziervogel et al. 2012) and deposited on the seafloor (Schrope 2013, Brooks et al. 2015). The rapid plume and settlement of hydrocarbon-plankton-bacterial product aggregation deposited in an event called the marine oil-snow sedimentation and flocculent accumulation (MOSSFA) (Brooks et al. 2015, Schwing et al. 2017b).

Consistent with these observations, in 2011, one year following the spill, rapid soluble and insoluble hydrocarbon deposition was detected in contaminated sediment in sites located in the DeSoto Canyon, including sites PCB06, XC2, and XC3 (Brooks et al. 2015, Romero et al. 2015) of the present study. Total PAH concentration of the sediment increased two to three fold (Romero et al. 2015). Sediments near the deeper plumes also experienced spikes in oil-degrading bacteria in September/October 2010 and in the summer seasons of 2012 – 2014 (Mason et al. 2014b, Overholt 2018).

Satellite measurements indicated surface plumes triggered a phytoplankton bloom over the canyon within weeks after the wellhead was capped (Hu et al. 2011). Elevated photosynthetic microbial groups in the top 1 cm of the sediment in November and December of 2010 also confirm the influence of the phytoplankton blooms (Brooks et al. 2015). Consistent with these observations, from 2010-2013, the sediment redoxcline sustained lasting changes indicative of an influx of enriched organic matter (Hastings et al. 2015). As a result of one or both of these perturbations, benthic foraminiferans in the canyon experienced a decline in density, species richness, and bioturbation overall of the sediment ceased, initially after the spill (Brooks et al. 2015, Schwing et al. 2015, Schwing et al. 2017a).

The distribution of highly depleted radiocarbon indicative of the DwH hydrocarbons were relatively light (Shantharam et al, in prep). Deposited hydrocarbons consisted of decayed, high molecular weight compounds *n*-alkanes (67%), low molecular weight *n*-alkanes (9%) and low weight PAHs (6%). This composition remained relatively unchanged for 3 years though large reductions in concentrations did occur for homohopanes (∼67%) and low weight compounds (*n*-alkanes and PAHs, ∼65% and ∼66% respectively) and to a lesser degree high molecular weight *n*-alkanes (∼43%) and PAHs (∼12%) (Romero et al. 2020). Perturbations to phytoplankton productivity largely abated by 2014 and 2015 (Li et al. 2019) over the canyon and sedimentary bacterial communities likely returned to baseline conditions (Yang et al. 2016, Liu et al. 2017). Between 2013 – 2016, sediment bioturbation resumed (Larson et al. 2018), redox steady-state conditions returned (Hastings et al. 2020), and foraminifera density and diversity increased and stabilized (Schwing et al. 2018, Schwing & Machain-Castillo 2020). Macrofauna for PCB06, XC2, XC3, S36, and XC4, in a similar timeframe (2012 – 2014) showed no change in richness and evenness, but elevated abundance in 2012 compared to 2013 and 2014 (Shantharam et al. In prep). Other macrofauna-based community stress and oil-impact indicators showed little to no signs of impact by 2014 and almost no difference from control sites by 2014 (Shantharam et al., in prep). Since the influence of oil at DeSoto Canyons sites seems to have tapered off by the 2014 sampling for the current study, the assumption is therefore made that the observed patterns are representative of the “typical” environmental forcing in the DeSoto Canyon region for sediment macrofauna, although potential exceptions are noted.

### 4.2 DeSoto Canyon macrofauna abundance, diversity, and community composition

Macrofauna in the DeSoto Canyon exhibited a general decrease in abundance with depth and between depth groups (Figure 2 and 4A respectively), consistent with some of the earliest GOM studies (Rowe & Menzel 1971, Rowe et al. 1974), previous deep-sea NGOM benthic faunal surveys and studies (Blake & Doyle 1983, Pequegnat et al. 1983, Pequegnat et al. 1990, Escobar-Briones et al. 1999), other studies of GOM canyons (Escobar-Briones et al. 2008) and the general deep sea (reviewed in Etter and Rex 2010). Peak abundance occurred at the shallowest stations at 485 m. This corresponds with earlier studies of northeastern GOM that reported max density between 355 and 650 m depending on season (Pequegnat et al. 1983, Pequegnat et al. 1990) and seems common to GOM macrofauna studies (Rowe & Menzel 1971, Rowe et al. 1974, Blake & Doyle 1983, Escobar-Briones et al. 1999, Stuart et al. 2016). Several studies also noted secondary peaks at around 1100 and 1500 m, in the central and western NGOM (Pequegnat et al. 1983, Pequegnat et al. 1990, Escobar-Briones et al. 1999, Stuart et al. 2016). In the current study these depths also have slightly higher values but not enough to stand out from the regression. Infaunal density in other large basins and depressions of the GOM report peak or high densities at similar depths. Baguley et al. (2006a) reported the highest density (9457 ind. m^-2^) for central NGOM meiofauna in the Mississippi Trough at 482 m.

The negative parabolic relationship observed for macrofaunal species richness with depth within DeSoto Canyon, with a peak at 1100 m, is comparable to the pattern observed for general NGOM fauna (Pequegnat et al. 1990, Haedrich et al. 2008, Stuart et al. 2016, Wei & Rowe 2019) and singular taxonomic groups over the larger GOM (Wicksten & Packard 2005, Reuscher & Shirley 2014, Shantharam & Baco 2019). Patterns of NGOM macrofauna richness are related to a host of environmental parameters that include food, habitat, pollution, and location (Haedrich et al. 2008), but the most influential, especially with depth, seems to be POC flux (Wei & Rowe 2019, Wei et al. accepted).

Evenness has not been reported in studies of NGOM macrofauna and only a few studies focused on canyons measure it. However, the classic increase of evenness with depth (Rex & Etter 2010) was observed within the Canyon and is consistent with what has been observed in the Scripps and La Jolla Canyons (∼0.30 – 0.80; Vetter and Dayton 1998), Nazaré Canyon (0.087 - 0.563; Curdia et al. 2004), the Whittard Canyon (0.662 – 0.923; Gunton 2015) and canyons of the Campos and Espirito Santo Basins off Brazil (∼0.58 – 0.90, Bernardino et al. 2019). Previous studies report a large range of evenness values in canyons, indicative of an inherent disturbance regime. Macrofaunal evenness in DeSoto Canyon is somewhat narrower than what has been reported in other canyons (0.7253 – 0.919) and although the Canyon has previously been described as inactive (Uchupi & Emery 1968, Bouma 1972), the range of evenness values reported here does not preclude an inherent disturbance regime. Cross-slope and deeper currents are known to be quite strong in the NGOM (Hamilton 1992, Hamilton & Lugo Fernandez L 2001) and can create a strong resonance in the narrowest part of the canyon at ∼715 m (Clarke & Van Gorder 2016) which theoretically may result in a flushing-type disturbance regime within Desoto akin to steeper-sided canyons.

### 4.3 DeSoto Canyon macrofauna composition, community structure, and association with environmental factors

Across the general NGOM Pequegnat et al (1990) first described three main depth zones for sediment macrofaunal assemblages: the Shelf/Slope-Transition (300 – 700 m), the Archibenthal Zone (700 – 1650 m), and the Abyssal (> 2000 m). Wei et al. (2010) had broader, overlapping depths with the NGOM divided into 4 zones, named the upper (213 – 542 m) with an extension submerging at 1572 m, mid and lower slope zones that split into eastern and western subzones (mid-eastern slope (625 – 1828 m), mid-western slope (863 – 1620 m), lower eastern slope (2275 – 3314 m), and lower western slope (2042 – 3008 m)), and also the abyssal plain (2954 - 3732). Within the DeSoto Canyon, macrofaunal community structure in this study showed three depth assemblages which largely fit into the regions of Pequegnat et al (1990); assemblage I at depths of 464 – 485 m, assemblage II at 669 – 1834 m, and assemblage III for sites greater than 2000 m.

Compositionally, the dominant macrofaunal groups maintained mostly similar proportions throughout the canyon, and the depth of peak abundances varied for all groups. Polychaetes dominated in the DeSoto Canyon, like most soft sediment continental margin environments (Gage & Tyler 1991, Grassle & Maciolek 1992), followed by crustaceans, and molluscs. While this coincides with previous NGOM surveys (Pequegnat et al. 1990), and some other Atlantic canyons (Gunton et al. 2015, Harriague et al. 2019), this pattern is not true of all canyons.

Polychaetes, while the most prevalent in submarine canyons in the Hawaiian islands, are followed by molluscs and then crustaceans are the next most common (De Leo et al. 2014). Hudson Canyon off New York state, also while dominated by polychaetes, has a strong proportion of bivalves, and sipunculans. (Rowe et al. 1982). Newport Canyon off California is strongly dominated by polychaetes, nemerteans, aplacophorans, and some echinoderms (Hartman 1963, Maurer et al. 1995). Adjacent canyons can show highly heterogeneous compositions as well. Cunha et al. (2011) report the Setúbal Canyon off Portugal has abundant taxa similar in proportion to the DeSoto Canyon but nearby Nazaré Canyon is predominated by molluscs, followed by polychaetes, arthropods, and echinoderms and the Cascais Canyon maintains crustaceans as the most abundant, then polychaetes, and then molluscs. The substrate can strongly determine the most abundant group in some canyons. Polychaetes and cumaceans, for example, are the most common in muddy/silty sections of the Carson Canyon off California, sandy sections had sipunculans and isopods, and the more gravel-heavy sections exhibited majority cumaceans and echinoderms (Houston & Haedrich 1984).

Within the DeSoto Canyon, some taxa, departed from the mean and had standout proportions at certain sites. Some of these disparate compositions may be attributable to hydrocarbon seep influence. Seeps occur in the canyon just as they do in with the larger GOM (MacDonald et al. 2015). Two sites sampled in the current study, Seep A and Peanut Mound, are known seeps, however since a video-guided multicorer was not employed, it could not be determined whether seep-influenced sediments were directly sampled or if general background sediments were sampled at these sites. Macrofauna in GOM seeps tends to consist of background GOM taxa and exhibit a large degree of heterogeneity in composition and community structure within seep microhabitat types (i.e., microbial mat, tubeworm, and soft sediment) (Washburn et al. 2018), typical of most seep habitats (Bernardino et al. 2012). Washburn et al. (2018) described NGOM seeps to generally be dominated by the polychaete Families Dorvilleidae, Hesionidae, and Ampharetidae, and DeSoto Canyon seeps sampled in the same study had high abundances of spionid and syllid polychaetes, and tanaid crustaceans. While none of the dominant GOM seep polychaete families were dominant in the samples of the current study, Seep A and Peanut Mound do show a high presence of syllids, spionids, and tanaids. Peanut Mound especially contained the most disparate community composition with the lowest proportion of polychaetes of the stations and the highest percentage of other crustaceans (isopods and cumaceans) and tanaids, though this did not yield a standout community structure in the NMDS. Cumaceans can especially be dominant on bacterial mats and sulfide seeps (Levin 2005). Thus these results may support the sampling of seeps at Seep A and Peanut Mound, however comparison to Washburn et al. (2018) is obfuscated by the coarse taxonomic resolution of the current data, the fact that syllids and spionids are some of the most diverse polychaete groups throughout the GOM (Reuscher & Shirley 2014), and that tanaids are generally dominant throughout the canyon sites sampled in the present study.

### 4.4 DeSoto Canyon wall vs. axis vs. the adjacent slope

The comparisons among the canyon axis, canyon wall and adjacent slope showed no difference in species richness or evenness among habitats, comparable to the findings of Wei and Rowe (2019). However, abundance was significantly higher on the canyon wall than the other habitats, and higher in the canyon axis compared to the adjacent slope. Higher abundance in the canyon is consistent with the high biomass previously observed in the canyon (Wei et al. 2012). The increased abundance on the canyon wall may be indicative of favorable environmental conditions. Parameters that were higher in at least one canyon habitat included salinity, oxygen, fluorescence, turbidity and percent sand. The parameters that showed the strongest correlation to community structure the DISTLM analyses were oxygen and POC flux. Greater oxygen in the canyon could overcome any limitation of the oxygen minimum zone observed in most mid-water regions of continental margins (Levin et al. 2001), however the lowest oxygen value of 4.22 mg/l would not be expected to be limiting to most macrofaunal species. Greater turbidity and fluorescence, potentially a product of a canyon-entrained water mass, would signify higher suspended particles in the canyon than outside and perhaps greater particulate organic matter flux to the sediments. Yet, this is contradicted by lower organic carbon and nitrogen in the canyon sediments compared to the adjacent slope. Average POC flux does show a trend of highest POC flux on the canyon wall, followed by the canyon axis, and lowest on the adjacent slope, but the differences were not statistically significant. This likely reflects the coarse resolution of the satellite measurements inadequately capturing habitat differences over a narrow geographic range but does not diminish the contribution that POC makes to community structure in general. Higher sediment organic matter has been found in the canyon before (Morse & Beazley 2008) and after (Brooks & Larson 2013, Chanton 2014) the DwH but did not appear to remain by 2014. Other unmeasured environmental variability may also influence canyon sediment macrofauna. The DeSoto Canyon contains a series of submarine channels, especially along the western wall, formed by mass movements that culminate in debris depots in the deeper basin (McAdoo et al. 2000). Sharp V-shaped incisions of these channels indicate high flushing and mass slumping until the channels reach the abyssal plain (Silva 2017). Additionally, strong currents occur along the narrow axis of the canyon (∼700 m) generated by subinertial canyon resonance (Clarke & Van Gorder 2016) can reach velocities to flush sediment in the canyon (A. Clarke, pers comm), and would potentially limit accumulation of organic material along the axis, explaining the lower abundance observed. Other canyon studies have also found sediment organic matter higher on the adjacent slope rather than the canyon as a result of high sedimental flushing (Liao et al. 2017).

### 4.5 Community structure across habitats

Based on the geographic locations of the bathymetric zones within the Gulf of Mexico designated by Wei et al. (2010), all sites from the current study, including the sites S35, S36, and S42 from Wei et al. (2010) that were revisited, should fall into the eastern mid-slope zone of that study. In Wei et al (2010), this zone included the DeSoto Canyon and extended east and west of the canyon with a slender portion reaching well into the western NGOM. However, the finer scale sampling of the current study revealed differences in community structure not only of the DeSoto Canyon sites from the adjacent slope, but also disparate structure of the canyon wall compared to the canyon axis.

Differences were also found between the environmental parameters tied to community structure. For the zones of Wei et al (2010), cluster analysis indicated that sites in this zone were highly influenced by POC mediated by the Mississippi River, dissolved oxygen, temperature, depth, sand, relative backscatter, and percent clay. In contrast, in the finer spatial scale of the current study, many of the grain size parameters, though significant individually, fell away in the DISTLM models and more emphasis was placed on oxygen and POC flux in the water column. In the summer, the season the canyon was sampled, cyclonic and anticyclonic eddies near the DeSoto Canyon can move low salinity, biologically productive Mississippi River output across the shelf and the head of the canyon (Müller Karger et al. 1991, Belabbassi et al. 2005, Walker L et al. 2005, Biggs et al. 2008, Jochens & DiMarco 2008). Strong thermohaline stratification prevents further intrusion into deeper waters, however (Jochens & DiMarco 2008). This leaves high salinity, highly oxygenated water characteristic of the North Atlantic Deepwater (NADW) (Rivas et al. 2005, Morse & Beazley 2008) to occupy the deep (>1000 m) sites, suggesting a strong influence of in situ seawater conditions on macrobenthic communities. Community structure differences between the habitat types may be driven in part by the higher abundance observed in the canyon, likewise the canyon wall over the canyon axis. Environmental factors that contribute to the difference in abundance among the habitats may also have an influence on community composition and community structure. These were discussed in the previous section. and included higher fluorescence, turbidity, and oxygen saturation in the canyon. The greater turbidity and fluorescence (measure of water-borne chlorophyll) could support a greater proportion of suspension-feeding bivalves and polychaetes and explain the higher abundances in the canyon. The difference in community structure also tracked with an eastward trend in longitude. Biggs et al. (2008) noted sea-surface chlorophyll was higher in the northeast GOM compared to the northwest, typically reaching a peak in the June-August timeframe and structuring the slope macrobenthos across the NGOM. This seasonality may also operate in smaller scale regions such as the DeSoto Canyon. Dissolved oxygen also demonstrates a longitudinal trend with decreasing values moving west to east but does not reach limiting levels and stands in contrast to the typical increase of oxygen at this depth (Jochens et al. 2005).

Differing community structure is not novel when comparing macrofaunal communities in canyons against the adjacent slope (Vetter & Dayton 1998, Duineveld et al. 2001, De Leo et al. 2014, Gunton et al. 2015, Bernardino et al. 2019a, Harriague et al. 2019). Disparate structures have been attributed to altered community composition that occurs because of topographical heterogeneity (De Leo et al. 2014) or higher organic loading in the canyon (Vetter & Dayton 1998, Duineveld et al. 2001, Gunton et al. 2015, Harriague et al. 2019). Many of the groups contributing to differences between DeSoto Canyon and the open slope communities were indicative of organic loading, such as thyasirid bivalves and opportunistic polychaetes.

Thyasirids especially are common in canyons where high organic deposition is present (Vetter & Dayton 1998, Cunha et al. 2011b, Bernardino et al. 2019a, Harriague et al. 2019) and can be the most discriminating taxon between canyon and adjacent slope habitats (Harriague et al. 2019).

While the difference in abundance and community structure between canyon and slope habitats is not unexpected, what drives differences within the canyon habitat communities remains elusive. Two sites which make up the canyon wall habitat, XC2 and XC3, in terms of taxonomic composition, contained proportions of molluscs higher than any of the other sites, with a high abundance of bivalves at XC3 and a high abundance of other molluscs at XC2. Noted seep-characteristic bivalve family Thyasiridae was several times more abundant at XC3 than other stations, contributing to that station’s high bivalve proportions. This hints at a chemosynthetic influence such as localized hydrocarbon seepage or bacterial decomposition of some type of organic enrichment, as has been observed in other canyons (Ingels et al. 2011, Bernardino et al. 2019a, Harriague et al. 2019). However, both XC2 and XC3 were also confirmed to have received DwH-induced sediment pulses and have been shown to have had high rates of organic matter respiration in the sediment following the spill (Hastings et al. 2015). Drawdowns in sediment porewater oxygen and the toxicity of petroleum aromatic hydrocarbons were responsible for an initial benthic decline (i.e., foraminiferans) (Schwing et al. 2015). But as the sediment environment recovered, the concomitant recolonization and succession of benthic fauna, along with any organic matter respiration, could have boosted benthic populations in XC2 and XC3. Thyasirids were found to be tolerant of DwH contamination (Washburn et al. 2016).

The organic enrichment observed (Hastings et al. 2015, Hastings et al. 2020) at XC3 may have bolstered bacterial production and provided an ideal habitat for this chemosymbiotic bivalve to expand in numbers. Thus, the higher macrofaunal abundance on the canyon wall in 2014 may be a remnant of that effect.

The confounding issue with either the seep or DwH argument is that sediment organic carbon was not particularly high for XC3 (average 1.54%). Differences might then instead be attributed to even smaller scale heterogeneity within the canyon. Channels along the wall exhibit a high amount of sinuosity and sediment accumulation along sediment channel curves (Silva 2017), potentially developing patches of high organic matter that could also explain the higher abundances observed. The greater sediment clay content on the canyon wall compared to the axis could reflect generally higher refractory organic matter content driving differences within the canyon. Further sampling at a higher resolution would not only better locate organically rich channel deposits but also enable identification of productive hydrocarbon seep habitats. It has been shown that there can be high turnover of canyon fauna on small spatial scales (< 100 m) (McClain & Barry 2010, Campanyà-Llovet et al. 2018) that can be driven by highly sporadic food patches (Campanyà-Llovet et al. 2018). The NGOM exhibits a high degree of microhabitat heterogeneity, on the order of centimeters to hundreds of kilometers, over singular isobaths (Nunnally et al. 2018) that seem to support the patch-mosaic model of Grassle and Sanders (1973). Thus, further research, at a finer sampling resolution, may be required to parse out the differences observed here.

## Acknowledgements

The authors would like to thank the captains and crew of the RV *Weatherbird* II and fellow cruise field PIs Ian MacDonald and Joel Kostka along with the many volunteers at sea. Numerous undergraduate and graduate sorters helped process sediment samples and identify specimens including Lauren Gillies-Campbell, Ben Labelle, Melissa Olguin, Kaitlin Hurley, Christine Palmer, Savannah Goode, Rose Luzader, Meaghan Fahletti, Chrissoula Rakowski, Andrea Schmidt, Morgan Harrison, Juliana De Andrade Souza, Madison Savage, Suhavi Kaur, Reena Manohar, Ashley Christine Alvarez, Travis Ferguson, Daniel Cardenas, Julie Andrews and Jefferson Hemphill. This research was made possible by a grant from the Gulf of Mexico Research Initiative to support the Deep-C: Deep Sea to Coast Connectivity in the Eastern Gulf of Mexico Research Consortium.

## References

Anderson M, Gorley RN, Clarke K (2008) PERMANOVA+ for primer: Guide to software and statistical methods

Anderson MJ, Walsh DC (2013) PERMANOVA, ANOSIM, and the Mantel test in the face of heterogeneous dispersions: What null hypothesis are you testing? Ecol Monogr 83:557–574

Antoine J, Bryant W (1968) The Major Transition Zones of the Gulf of Mexico: Desoto and Campeche Canyons. AAPG Bull 52:1831–1831

Arzola RG, Wynn RB, Lastras G, Masson DG, Weaver PP (2008) Sedimentary features and processes in the Nazaré and Setúbal submarine canyons, west Iberian margin. Mar Geol 250:64–88

Baguley JG, Montagna PA, Hyde LJ, Kalke RD, Rowe GT (2006a) Metazoan meiofauna abundance in relation to environmental variables in the northern Gulf of Mexico deep sea. Deep Sea Research Part I: Oceanographic Research Papers 53:1344–1362

Baguley JG, Montagna PA, Lee W, Hyde LJ, Rowe GT (2006b) Spatial and bathymetric trends in Harpacticoida (Copepoda) community structure in the Northern Gulf of Mexico deep-sea. J Exp Mar Biol Ecol 330:327–341

Balsam WL, Beeson JP (2003) Sea-floor sediment distribution in the Gulf of Mexico. Deep Sea Research Part I: Oceanographic Research Papers 50:1421–1444

Behrenfeld MJ, Falkowski PG (1997) Photosynthetic rates derived from satellite based chlorophyll concentration. Limnol Oceanogr 42:1–20 based chlorophyll

Belabbassi L, Chapman P, Nowlin Jr WD, Jochens AE, Biggs DC (2005) Summertime nutrient supply to near-surface waters of the northeastern Gulf of Mexico: 1998, 1999, and 2000. Gulf Mex Sci 23:1

Bernardino AF, Gama RN, Mazzuco ACA, Omena EP, Lavrado HP (2019a) Submarine canyons support distinct macrofaunal assemblages on the deep SE Brazil margin. Deep Sea Research Part I: Oceanographic Research Papers 149:103052

Bernardino AF, Gama RN, Mazzuco ACA, Omena EP, Lavrado HP (2019b) Submarine canyons support distinct macrofaunal assemblages on the deep SE Brazil margin. Deep Sea Research Part I: Oceanographic Research Papers

Bernardino AF, Levin LA, Thurber AR, Smith CR (2012) Comparative Composition, Diversity and Trophic Ecology of Sediment Macrofauna at Vents, Seeps and Organic Falls. PLOS ONE 7:e33515

Biggs DC, Hu C, Müller-Karger FE (2008) Remotely sensed sea-surface chlorophyll and POC flux at Deep Gulf of Mexico Benthos sampling stations. Deep Sea Research Part II: Topical Studies in Oceanography 55:2555–2562

Blake NJ, Doyle LJ (1983) Infaunal-sediment relationships at the shelf-slope break. SEPM Special Publication:381–389

Bouma AH (1972) Distribution of sediments and sedimentary structures in the Gulf of Mexico.

Brodeur RD (2001) Habitat-specific distribution of Pacific ocean perch (Sebastes alutus) in Pribilof Canyon, Bering Sea. Cont Shelf Res 21:207–224

Brooks GR, Larson RA (2013) Sediment texture and composition, NE Gulf of Mexico, 2010-2013. Gulf of Mexico Research Initiative Information and Data Cooperative (GRIIDC), Corpus Christi, TX

Brooks GR, Larson RA, Schwing PT, Romero I, Moore C, Reichart G-J, Jilbert T, Chanton JP, Hastings DW, Overholt WA (2015) Sedimentation pulse in the NE Gulf of Mexico following the 2010 DWH blowout. PLOS ONE 10:e0132341

Burnham KP, Anderson DR (2004) Multimodel inference: understanding AIC and BIC in model selection. Sociol Methods Res 33:261–304

Burrough PA, McDonnell R, McDonnell RA, Lloyd CD (2015) Principles of geographical information systems. Oxford university press

Camilli R, Reddy CM, Yoerger DR, Van Mooy BA, Jakuba MV, Kinsey JC, McIntyre CP, Sylva SP, Maloney JV (2010) Tracking hydrocarbon plume transport and biodegradation at Deepwater Horizon. Science 330:201–204

Campanyà-Llovet N, Snelgrove PVR, De Leo FC (2018) Food quantity and quality in Barkley Canyon (NE Pacific) and its influence on macroinfaunal community structure. Prog Oceanogr 169:106–119

Canals M, Puig P, de Madron XD, Heussner S, Palanques A, Fabres J (2006) Flushing submarine canyons. Nature 444:354–357

Chanton J, Zhao T, Rosenheim BE, Joye SB, Bosman S, Brunner CA, Yeager KM, Diercks AR, Hollander D (2014) Using Natural Abundance Radiocarbon to trace the Flux of Petrocarbon to the Seafloor following the Deepwater Horizon Oil Spill. Environ Sci Technol 49:847–854

Chanton JP (2014) Radiocarbon measurements on surface sediment organic matter following the Deepwater Horizon Oil Spill, 2010-2012. Gulf of Mexico Research Initiative Information and Data Cooperative (GRIIDC), Corpus Christi, TX

Clarke AJ, Van Gorder S (2016) Subinertial canyon resonance. Geophys Res Lett 43:3872–3879

Clarke K, Gorley R (2015) PRIMER v7: User Manual/Tutorial; PRIMER-E: Plymouth, UK, 2015. PRIMER-E, Plymouth, UK

Coleman FC, Chanton JP, Chassignet EP (2014) Ecological Connectivity in Northeastern Gulf of Mexico – The Deep-C Initiative. International Oil Spill Conference Proceedings 2014:1972–1984

Company JB, Puig P, Sarda F, Palanques A, Latasa M, Scharek R (2008) Climate influence on deep sea populations. PLOS ONE 3:e1431

Cunha MR, Paterson GL, Amaro T, Blackbird S, de Stigter HC, Ferreira C, Glover A, Hilario A, Kiriakoulakis K, Neal L (2011a) Biodiversity of macrofaunal assemblages from three Portuguese submarine canyons (NE Atlantic). Deep Sea Research Part II: Topical Studies in Oceanography 58:2433–2447

Cunha MR, Paterson GLJ, Amaro T, Blackbird S, de Stigter HC, Ferreira C, Glover A, Hilário A, Kiriakoulakis K, Neal L, Ravara A, Rodrigues CF, Tiago Á, Billett DSM (2011b) Biodiversity of macrofaunal assemblages from three Portuguese submarine canyons (NE Atlantic). Deep Sea Research Part II: Topical Studies in Oceanography 58:2433–2447

Curdia J, Carvalho S, Ravara A, Gage J, Rodrigues A, Quintino V (2004) Deep macrobenthic communities from Nazaré submarine canyon (NW Portugal). Sci Mar 68:171–180

De Leo FC, Drazen JC, Vetter EW, Rowden AA, Smith CR (2012) The effects of submarine canyons and the oxygen minimum zone on deep-sea fish assemblages off Hawai’i. Deep Sea Research Part I: Oceanographic Research Papers 64:54–70

De Leo FC, Smith CR, Rowden AA, Bowden DA, Clark MR (2010) Submarine canyons: hotspots of benthic biomass and productivity in the deep sea. Proceedings of the Royal Society of London B: Biological Sciences:rspb20100462

De Leo FC, Vetter EW, Smith CR, Rowden AA, McGranaghan M (2014) Spatial scale-dependent habitat heterogeneity influences submarine canyon macrofaunal abundance and diversity off the Main and Northwest Hawaiian Islands. Deep Sea Research Part II: Topical Studies in Oceanography 104:267–290

de Stigter HC, Boer W, de Jesus Mendes PA, Jesus CC, Thomsen L, van den Bergh GD, van Weering TC (2007) Recent sediment transport and deposition in the Nazaré Canyon, Portuguese continental margin. Mar Geol 246:144–164

Doyle LJ, Sparks TN (1980) Sediments of the Mississippi, Alabama, and Florida (MAFLA) continental shelf. J Sediment Res 50:905–915

Duineveld G, Lavaleye M, Berghuis E, de Wilde P (2001) Activity and composition of the benthic fauna in the Whittard Canyon and the adjacent continental slope (NE Atlantic). Oceanol Acta 24:69–83

Ebbe B, Billett DS, Brandt A, Ellingsen K, Glover A, Keller S, Malyutina M, Martínez Arbizu P, Molodtsova T, Rex M (2010) Diversity of abyssal marine life. Life in the World’s Oceans: Diversity, Distribution, and Abundance, edited by: McIntyre, A:139–160

Emiliani C, Gartner S, Lidz B, Eldridge K, Elvey DK, Huang TC, Stipp JJ, Swanson MF (1975) Paleoclimatological analysis of late Quaternary cores from the northeastern Gulf of Mexico. Science 189:1083–1088

Escobar-Briones E, Santillán ELE, Legendre P (2008) Macrofaunal density and biomass in the Campeche Canyon, Southwestern Gulf of Mexico. Deep Sea Research Part II: Topical Studies in Oceanography 55:2679–2685

Escobar-Briones E, Signoret M, Hernández D (1999) Variation of the macrobenthic infaunal density in a bathymetric gradient: Western Gulf of Mexico. Cienc Mar 25:193–212

Gage JD (1996) Why are there so many species in deep-sea sediments? J Exp Mar Biol Ecol 200:257–286

Gage JD, Tyler PA (1991) Deep-sea biology: a natural history of organisms at the deep-sea floor.Cambridge University Press

Garcia-Pineda O, Macdonald I, Hu C, Svejkovsky J, Hess M, Dukhovskoy D, Morey SL (2013) Detection of floating oil anomalies from the Deepwater Horizon oil spill with synthetic aperture radar. Oceanography 26:124–137

Genin A (2004) Bio-physical coupling in the formation of zooplankton and fish aggregations over abrupt topographies. J Mar Syst 50:3–20

Gerino M, Stora G, Poydenot F, Bourcier M (1995) Benthic fauna and bioturbation on the Mediterranean continental slope: Toulon Canyon. Cont Shelf Res 15:1483–1496

Gesteira JG, Dauvin J, Fraga MS (2003) Taxonomic level for assessing oil spill effects on soft-bottom sublittoral benthic communities. Mar Pollut Bull 46:562–572

Gould HR, Stewart RH (1955) Continental terrace sediments in the northeastern Gulf of Mexico. Special Publications of SEPM Finding Ancient Shorelines:2–20

Grassle JF, Maciolek NJ (1992) Deep-sea species richness: regional and local diversity estimates from quantitative bottom samples. Am Nat:313–341

Grassle JF, Sanders HL Life histories and the role of disturbance. Proc Deep Sea Research and Oceanographic Abstracts. Elsevier

Greene C, Wiebe P, Burczynski J, Youngbluth M (1988) Acoustical detection of high-density krill demersal layers in the submarine canyons off Georges Bank. Science 241:359–361

Gunton L (2015) Deep-sea macrofaunal biodiversity of the Whittard Canyon (NE Atlantic). PhD, University of Southampton,

Gunton LM, Gooday AJ, Glover AG, Bett BJ (2015) Macrofaunal abundance and community composition at lower bathyal depths in different branches of the Whittard Canyon and on the adjacent slope (3500 m; NE Atlantic). Deep Sea Research Part I: Oceanographic Research Papers 97:29–39

Haedrich RL, Devine JA, Kendall VJ (2008) Predictors of species richness in the deep-benthic fauna of the northern Gulf of Mexico. Deep Sea Research Part II: Topical Studies in Oceanography 55:2650–2656

Hamilton P (1992) Lower continental slope cyclonic eddies in the central Gulf of Mexico. Journal of Geophysical Research: Oceans 97:2185–2200

Hamilton P, Lugo D Fernandez A (2001) Observations of high speed deep currents in the northern Gulf of Mexico. Geophys Res Lett 28:2867–2870

Hamilton P, Speer K, Snyder R, Wienders N, Leben RR (2015) Shelf break exchange events near the De Soto Canyon. Cont Shelf Res 110:25–38

Harriague AC, Danovaro R, Misic C (2019) Macrofaunal assemblages in canyon and adjacent slope of the NW and Central Mediterranean systems. Prog Oceanogr 171:38–48

Harris P, Macmillan-Lawler M, Rupp J, Baker E (2014) Geomorphology of the oceans. Mar Geol 352:4–24

Harris PT, Whiteway T (2011) Global distribution of large submarine canyons: Geomorphic differences between active and passive continental margins. Mar Geol 285:69–86

Harrold C, Light K, Lisin S (2003) Organic enrichment of submarine canyon and continental shelf benthic communities by macroalgal drift imported from nearshore kelp forests. Limnol Oceanogr 43:669–678

Hartman O (1963) Quantitative survey of the benthos of San Pedro Basin, Southern California. Part II Biology. Allan Hancock Pacific Expedition 27:424

Hastings DW, Bartlett T, Brooks GR, Larson RA, Quinn KA, Razionale D, Schwing PT, Bernal LHP, Ruiz-Fernández AC, Sánchez-Cabeza J-A (2020) Changes in Redox Conditions of Surface Sediments Following the Deepwater Horizon and Ixtoc 1 Events. Deep Oil Spills. Springer

Hastings DW, Schwing PT, Brooks GR, Larson RA, Morford JL, Roeder T, Quinn KA, Bartlett T, Romero IC, Hollander DJ (2015) Changes in sediment redox conditions following the BP DWH blowout event. Deep Sea Research Part II: Topical Studies in Oceanography 129:167–198

Hazen TC, Dubinsky EA, DeSantis TZ, Andersen GL, Piceno YM, Singh N, Jansson JK, Probst A, Borglin SE, Fortney JL (2010) Deep-sea oil plume enriches indigenous oil-degrading bacteria. Science 330:204–208

Hickey BM (1997) The response of a steep-sided, narrow canyon to time-variable wind forcing. J Phys Oceanogr 27:697–726

Houston K, Haedrich R (1984) Abundance and biomass of macrobenthos in the vicinity of Carson Submarine Canyon, northwest Atlantic Ocean. Mar Biol 82:301–305

Hu C, Weisberg RH, Liu Y, Zheng L, Daly KL, English DC, Zhao J, Vargo GA (2011) Did the northeastern Gulf of Mexico become greener after the Deepwater Horizon oil spill? Geophys Res Lett 38

Hunter W, Jamieson A, Huvenne V, Witte U (2013) Sediment community responses to marine vs. terrigenous organic matter in a submarine canyon. Biogeosciences 10:67–80

Ingels J, Tchesunov AV, Vanreusel A (2011) Meiofauna in the Gollum Channels and the Whittard Canyon, Celtic Margin—how local environmental conditions shape nematode structure and function. PLOS ONE 6:e20094

Jackson MLR (1969) Soil Chemical Analysis: Advanced Course : a Manual of Methods Useful for Instruction and Research in Soil Chemistry, Physical Chemistry of Soils, Soil Fertility, and Soil Genesis. M.L. Jackson

Jochens AE, Bender L, DiMarco S, Morse J, Kennicutt MC, Howard M, Nowlin Jr WD (2005) Understanding the Processes that Maintain the Oxygen Levels in the Deep Gulf of Mexico: Synthesis Report. In: Interior UDot (ed). Bureau of Ocean Energy Management

Jochens AE, DiMarco SF (2008) Physical oceanographic conditions in the deepwater Gulf of Mexico in summer 2000–2002. Deep Sea Research Part II: Topical Studies in Oceanography 55:2541–2554

Kessler JD, Valentine DL, Redmond MC, Du M, Chan EW, Mendes SD, Quiroz EW, Villanueva CJ, Shusta SS, Werra LM (2011) A persistent oxygen anomaly reveals the fate of spilled methane in the deep Gulf of Mexico. Science 331:312–315

Kleindienst S, Grim S, Sogin M, Bracco A, Crespo-Medina M, Joye SB (2015) Diverse, rare microbial taxa responded to the Deepwater Horizon deep-sea hydrocarbon plume. ISME J

Klinck JM (1996) Circulation near submarine canyons: A modeling study. Journal of Geophysical Research: Oceans 101:1211–1223

Konert M, Vandenberghe J (1997) Comparison of laser grain size analysis with pipette and sieve analysis: a solution for the underestimation of the clay fraction. Sedimentology 44:523–535

Larson RA, Brooks GR, Schwing PT, Holmes CW, Carter SR, Hollander DJ (2018) High-resolution investigation of event driven sedimentation: Northeastern Gulf of Mexico. Anthropocene 24:40–50

Lavoie D, Simard Y, Saucier FJ (2000) Aggregation and dispersion of krill at channel heads and shelf edges: the dynamics in the Saguenay-St. Lawrence Marine Park. Can J Fish Aquat Sci 57:1853–1869

Levin L, Etter R, Rex M, Gooday A, Smith C, Pineda J, Stuart CT, Hessler R, Pawson D (2001) Environmental Influences on Regional Deep-Sea Species Diversity. Annu Rev Ecol Syst 32:51–93

Levin LA (2005) Ecology of cold seep sediments: Interactions of fauna with flow, chemistry and microbes. Oceanogr Mar Biol 43:1–46

Li Y, Hu C, Quigg A, Gao H (2019) Potential influence of the Deepwater Horizon oil spill on phytoplankton primary productivity in the northern Gulf of Mexico. Environ Res Lett 14:094018

Liao J-X, Chen G-M, Chiou M-D, Jan S, Wei C-L (2017) Internal tides affect benthic community structure in an energetic submarine canyon off SW Taiwan. Deep Sea Research Part I: Oceanographic Research Papers 125:147–160

Liu J, Bacosa HP, Liu Z (2017) Potential environmental factors affecting oil-degrading bacterial populations in deep and surface waters of the northern Gulf of Mexico. Front Microbiol 7:2131

Lutz MJ, Caldeira K, Dunbar RB, Behrenfeld MJ (2007) Seasonal rhythms of net primary production and particulate organic carbon flux to depth describe the efficiency of biological pump in the global ocean. Journal of Geophysical Research: Oceans (1978–2012) 112:C10011

MacDonald IR, Garcia-Pineda O, Beet A, Daneshgar Asl S, Feng L, Graettinger G, French-McCay D, Holmes J, Hu C, Huffer F, Leifer I, Muller-Karger F, Solow A, Silva M, Swayze G (2015) Natural and unnatural oil slicks in the Gulf of Mexico. Journal of Geophysical Research: Oceans 120:8364–8380

Mason O, Han J, Woyke T, Jansson J (2014a) Single-cell genomics reveals features of a *Colwellia* species that was dominant during the Deepwater Horizon oil spill. Front Microbiol 5:332

Mason OU, Scott NM, Gonzalez A, Robbins-Pianka A, Bælum J, Kimbrel J, Bouskill NJ, Prestat E, Borglin S, Joyner DC, Fortney J, Jurelevicius D, Stringfellow WT, Hazen TC, Knight R, Gilbert JA, Jansson JK (2014b) Metagenomics reveals sediment microbial community response to Deepwater Horizon oil spill. ISME J 8:1464–1475

Maurer D, Robertson G, Gerlinger T (1995) Community Structure of Soft Bottom Macrobenthos of the Newport Submarine Canyon, California. Marine Ecology 16:57–72

McAdoo B, Pratson L, Orange D (2000) Submarine landslide geomorphology, US continental slope. Mar Geol 169:103–136

McClain C, R., Nunnally C, Benfield Mark C (2019) Persistent and substantial impacts of the Deepwater Horizon oil spill on deep-sea megafauna. Royal Society Open Science 6:191164

McClain CR, Barry JP (2010) Habitat heterogeneity, disturbance, and productivity work in concert to regulate biodiversity in deep submarine canyons. Ecology 91:964–976

McNutt MK, Camilli R, Crone TJ, Guthrie GD, Hsieh PA, Ryerson TB, Savas O, Shaffer F (2012) Review of flow rate estimates of the Deepwater Horizon oil spill. Proc Natl Acad Sci USA 109:20260–20267

Morse JW, Beazley MJ (2008) Organic matter in deepwater sediments of the Northern Gulf of Mexico and its relationship to the distribution of benthic organisms. Deep Sea Research Part II: Topical Studies in Oceanography 55:2563–2571

Müller D Karger FE, Walsh JJ, Evans RH, Meyers MB (1991) On the seasonal phytoplankton concentration and sea surface temperature cycles of the Gulf of Mexico as determined by satellites. Journal of Geophysical Research: Oceans 96:12645–12665

Nunnally C, Landry C, Gholson S, McClain C (2018) Patchiness of sediment communities in the deep Gulf of Mexico across several spatial scales indicate a diversity of microscale habitat differences that drive diversity. Ocean Science Meeting, Portland, OR

Nürnberg D, Ziegler M, Karas C, Tiedemann R, Schmidt MW (2008) Interacting Loop Current variability and Mississippi River discharge over the past 400 kyr. Earth Planet Sci Lett 272:278–289

Oliveira A, Santos A, Rodrigues A, Vitorino J (2007) Sedimentary particle distribution and dynamics on the Nazaré canyon system and adjacent shelf (Portugal). Mar Geol 246:105–122

Overholt WA (2018) The response of marine benthic microbial populations to the Deepwater Horizon oil spill. Ph.D., Georgia Institute of Technology, Atlanta, GA

Passow U, Ziervogel K, Asper V, Diercks A (2012) Marine snow formation in the aftermath of the Deepwater Horizon oil spill in the Gulf of Mexico. Environmental Research Letters 7:035301

Paterson GL, Glover AG, Cunha MR, Neal L, de Stigter HC, Kiriakoulakis K, Billett DS, Wolff GA, Tiago A, Ravara A (2011) Disturbance, productivity and diversity in deep-sea canyons: A worm’s eye view. Deep Sea Research Part II: Topical Studies in Oceanography 58:2448–2460

Pequegnat W, Pequegnat L, Kleypas J, James B, Kennedy E, Hubbard G (1983) The ecological communities of the continental slope and adjacent regimes of the northern Gulf of Mexico. Final report to US Dept. of the Interior, Minerals Management Service, Gulf of Mexico OCS Region, New Orleans, LA.(Contract No. AA851-CT1-12)

Pequegnat WE, Gallaway BJ, Pequegnat LH (1990) Aspects of the ecology of the deep-water fauna of the Gulf of Mexico. Am Zool 30:45–64

Redmond MC, Valentine DL (2012) Natural gas and temperature structured a microbial community response to the Deepwater Horizon oil spill. Proceedings of the National Academy of Sciences 109:20292–20297

Reuscher MG, Shirley TC (2014) Diversity, distribution, and zoogeography of benthic polychaetes in the Gulf of Mexico. Marine Biodiversity 44:519–532

Rex MA, Etter RJ (2010) Deep-sea biodiversity: pattern and scale. Harvard University Press

Rinehart R, Wright DJ, Lundblad ER, Larkin EM, Murphy J, Cary-Kothera L ArcGIS 8. x benthic terrain modeler: Analysis in American Samoa. Proc Proceedings of the 24th Annual ESRI User Conference, San Diego, CA

Rivas D, Badan A, Ochoa J (2005) The Ventilation of the Deep Gulf of Mexico. J Phys Oceanogr 35:1763–1781

Romero I, Schwing P, Brooks G, Larson R, Hastings D, Ellis G, Goddard E, Hollander D (2015) Hydrocarbons in Deep-Sea Sediments following the 2010 Deepwater Horizon Blowout in the Northeast Gulf of Mexico. PLOS ONE 10:e0128371–e0128371

Romero IC, Chanton JP, Roseheim BE, Radović JR, Schwing PT, Hollander DJ, Larter SR, Oldenburg TB (2020) Long-Term Preservation of Oil Spill Events in Sediments: The Case for the Deepwater Horizon Oil Spill in the Northern Gulf of Mexico. Deep Oil Spills. Springer

Rowe GT, Menzel DW (1971) Quantitative benthic samples from the deep Gulf of Mexico with some comments on the measurement of deep-sea biomass. Bull Mar Sci 21:556–566

Rowe GT, Morse J, Nunnally C, Boland GS (2008) Sediment community oxygen consumption in the deep Gulf of Mexico. Deep Sea Research Part II: Topical Studies in Oceanography 55:2686–2691

Rowe GT, Polloni PT, Haedrich RL (1982) The deep-sea macrobenthos on the continental margin of the northwest Atlantic Ocean. Deep Sea Research Part A. Oceanographic Research Papers 29:257–278

Rowe GT, Polloni PT, Horner S Benthic biomass estimates from the northwestern Atlantic Ocean and the northern Gulf of Mexico. Proc Deep Sea Research and Oceanographic Abstracts. Elsevier

Schlacher TA, Schlacher-Hoenlinger M, A., Williams A, Althaus F, Hooper J, Kloser R (2007) Richness and distribution of sponge megabenthos in continental margin canyons off southeastern Australia. Mar Ecol Prog Ser 340:73–88

Schrope M (2013) Dirty blizzard buried Deepwater Horizon oil. Nature

Schwing P, O’malley B, Romero I, Martínez-Colón M, Hastings D, Glabach M, Hladky E, Greco A, Hollander D (2017a) Characterizing the variability of benthic foraminifera in the northeastern Gulf of Mexico following the Deepwater Horizon event (2010–2012). Environmental Science and Pollution Research 24:2754–2769

Schwing PT, Brooks GR, Larson R, Holmes C, O’Malley B, Hollander DJ (2017b) Constraining the spatial extent of Marine Oil Snow Sedimentation and Flocculent Accumulation (MOSSFA) following the Deepwater Horizon Event using an excess 210Pb flux approach. Environ Sci Technol 51:5962–5968

Schwing PT, Machain-Castillo ML (2020) Impact and Resilience of Benthic Foraminifera in the Aftermath of the Deepwater Horizon and Ixtoc 1 Oil Spills. Deep Oil Spills. Springer

Schwing PT, O’Malley BJ, Hollander DJ (2018) Resilience of benthic foraminifera in the Northern Gulf of Mexico following the Deepwater Horizon event (2011–2015). Ecol Indic 84:753–764

Schwing PT, Romero IC, Brooks GR, Hastings DW, Larson RA, Hollander DJ (2015) A Decline in Benthic Foraminifera following the Deepwater Horizon Event in the Northeastern Gulf of Mexico. PLOS ONE 10:e0120565

Shantharam AK, Baco AR (2019) Biogeographic and Bathymetric Patterns of Benthic Molluscs in the Gulf of Mexico. Deep Sea Research Part I: Oceanographic Research Papers:103167

Shantharam AK, Wei C-L, Baco A (In prep) Interannual temporal patterns of DeSoto Canyon macrofauna and resilience to the Deepwater Horizon Oil Spill. Mar Pollut Bull

Silva MG (2017) Fate of the Mesophotic Coral Ecosystem (MCE) in the Northeastern Gulf of Mexico after the Deepwater Horizon Incident. Ph.D., Florida State University, Tallahassee, FL

Snelgrove PV (1999) Getting to the bottom of marine biodiversity: sedimentary habitats: ocean bottoms are the most widespread habitat on earth and support high biodiversity and key ecosystem services. Bioscience 49:129–138

Socolofsky SA, Adams EE, Sherwood CR (2011) Formation dynamics of subsurface hydrocarbon intrusions following the Deepwater Horizon blowout. Geophys Res Lett 38:L09602

Somerfield P, Clarke K (1995) Taxonomic levels, in marine community studies, revisited. Marine ecology progress series. Oldendorf 127:113–119

Sorbe JC (1999) Deep-sea macrofaunal assemblages within the benthic boundary layer of the Cap-Ferret Canyon (Bay of Biscay, NE Atlantic). Deep Sea Research Part II: Topical Studies in Oceanography 46:2309–2329

Stuart CT, Brault S, Rowe GT, Wei CL, Wagstaff M, McClain CR, Rex MA (2016) Nestedness and species replacement along bathymetric gradients in the deep sea reflect productivity: a test with polychaete assemblages in the oligotrophic north west Gulf of Mexico. J Biogeogr 44:548–555

Team RC (2019) R: A language and environment for statistical computing. R Foundation for Statistical Computing, Vienna, Austria

Tyler P, Amaro T, Arzola R, Cunha MR, De Stigter H, Gooday A, Huvenne V, Ingels J, Kiriakoulakis K, Lastras G, Masson D, Oliveira A, Pattenden A, Vanreusel ANN, Van Weering T, Vitorino J, Witte U, Wolff G (2009) Europe’s Grand Canyon Nazare Submarine Canyon. Oceanography 22:46–57

Uchupi E, Emery KO (1968) Structure of continental margin off Gulf Coast of United States. AAPG Bull 52:1162–1193

Uiblein F, Lorance P, Latrouite D (2003) Behaviour and habitat utilisation of seven demersal fish species on the Bay of Biscay continental slope, NE Atlantic. Mar Ecol Prog Ser 257:223–232

Valentine DL, Fisher GB, Bagby SC, Nelson RK, Reddy CM, Sylva SP, Woo MA (2014) Fallout plume of submerged oil from Deepwater Horizon. Proc Natl Acad Sci USA 111:15906–15911

Valentine DL, Kessler JD, Redmond MC, Mendes SD, Heintz MB, Farwell C, Hu L, Kinnaman FS, Yvon-Lewis S, Du M (2010) Propane respiration jump-starts microbial response to a deep oil spill. Science 330:208–211

Vetter E, Dayton P (1998) Macrofaunal communities within and adjacent to a detritus-rich submarine canyon system. Deep Sea Research Part II: Topical Studies in Oceanography 45:25–54

Vetter E, Dayton P (1999) Organic enrichment by macrophyte detritus, and abundance patterns of megafaunal populations in submarine canyons. Marine ecology. Progress series 186:137–148

Vetter EW (1994) Hotspots of benthic production. Nature 372:47

Vetter EW, Smith CR, De Leo FC (2010) Hawaiian hotspots: enhanced megafaunal abundance and diversity in submarine canyons on the oceanic islands of Hawaii. Marine Ecology 31:183–199

Walker ND, Wiseman Jr WJ, Rouse Jr LJ, Babin A (2005) Effects of river discharge, wind stress, and slope eddies on circulation and the satellite-observed structure of the Mississippi River plume. J Coast Res:1228–1244

Warwick R (1988) Analysis of community attributes of the macrobenthos of Frierfjord/Langesundfjord at taxonomic levels higher than species. Mar Ecol Prog Ser 46:167–170

Washburn T, Rhodes AC, Montagna PA (2016) Benthic taxa as potential indicators of a deep-sea oil spill. Ecol Indic 71:587–597

Washburn TW, Demopoulos AWJ, Montagna PA (2018) Macrobenthic infaunal communities associated with deep-sea hydrocarbon seeps in the northern Gulf of Mexico. Marine Ecology 39:e12508

Wei C-L, Chen M, Wicksten MK, Rowe Gilbert T (accepted) Macrofauna bivalve diversity from the deep northern Gulf of Mexico. Ecol Res

Wei C-L, Rowe GT (2019) Productivity controls macrofauna diversity in the deep northern Gulf of Mexico. Deep Sea Research Part I: Oceanographic Research Papers 143:17–27

Wei C-L, Rowe GT, Escobar-Briones E, Nunnally C, Soliman Y, Ellis N (2012) Standing Stocks and Body Size of Deep-sea Macrofauna: Predicting the Baseline of 2010 Deepwater Horizon Oil Spill in the Northern Gulf of Mexico. Deep Sea Research Part I: Oceanographic Research Papers

Wei C-L, Rowe GT, Hubbard GF, Scheltema AH, Wilson GD, Petrescu I, Foster JM, Wicksten MK, Chen M, Davenport R, Soliman YS, Wang Y (2010) Bathymetric zonation of deep-sea macrofauna in relation to export of surface phytoplankton production. Mar Ecol Prog Ser 399:1–14

Wicksten MK, Packard JM (2005) A qualitative zoogeographic analysis of decapod crustaceans of the continental slopes and abyssal plain of the Gulf of Mexico. Deep Sea Research Part I: Oceanographic Research Papers 52:1745–1765

Yang T, Nigro LM, Gutierrez T, Joye SB, Highsmith R, Teske A (2016) Pulsed blooms and persistent oil-degrading bacterial populations in the water column during and after the Deepwater Horizon blowout. Deep Sea Research Part II: Topical Studies in Oceanography 129:282–291

Yeager KM, Santschi PH, Rowe GT (2004) Sediment accumulation and radionuclide inventories ^239,240^Pu, ^210^Pb and ^234^Th) in the northern Gulf of Mexico, as influenced by organic matter and macrofaunal density. Mar Chem 91:1–14

Yoklavich MM, Greene HG, Cailliet GM, Sullivan DE, Lea RN, Love MS (2000) Habitat associations of deep-water rockfishes in a submarine canyon: an example of a natural refuge. Fishery Bulletin-National Oceanic and Atmospheric Administration 98:625–641

Ziervogel K, McKay L, Rhodes B, Osburn CL, Dickson-Brown J, Arnosti C, Teske A (2012) Microbial activities and dissolved organic matter dynamics in oil-contaminated surface seawater from the Deepwater Horizon oil spill site. PLOS ONE 7:e34816

